# Assessment of cortical excitability in awake rhesus macaques with transcranial magnetic stimulation: translational insights from recruitment curves

**DOI:** 10.1101/2024.12.17.628832

**Authors:** Anna Padányi, Balázs Knakker, Balázs Lendvai, István Hernádi

## Abstract

**Background and objectives:** Cortical excitability (CE) is commonly assessed by recording motor evoked potentials (MEPs) in response to single-pulse transcranial magnetic stimulation (sp-TMS). While the motor threshold (MT) remains the most widely used measure of CE, it provides a one-dimensional, criterion-based assessment. In contrast, the recruitment curve (RC) offers a more comprehensive characterization of the full dynamics of cortical recruitment. Yet, only a few preclinical studies involving translationally relevant non-human primates were conducted, and most were under anaesthesia. Hence, we aimed to characterise CE in awake rhesus macaques by recording traditionally defined MT and RCs.

**Methods:** Traditional MT with a 100 µV MEP criterion (‘tradMT’) was measured in 8 awake adult male rhesus macaques using C-B65 coil and MagVenture stimulator. RCs were recorded at nine relative intensity levels (0.5 – 1.5 × tradMT) in 4 macaques. A sigmoid function was fitted to obtain key CE parameters: the inflection point, lower ankle point, and plateau.

**Results:** TradMT values were stable and replicable, and aligned most closely with the inflection point of the RC. The lower ankle points were found around at 0.9 × tradMT, marking the transition from a constant to a logarithmic phase, representing a physiologically relevant threshold. Plateau MEP amplitudes were substantially smaller compared to those reported in humans.

**Conclusion:** Fitted RC parameters revealed a distinction between tradMT and the physiologically relevant threshold. The overall RC shape was consistent with human data, suggesting similar recruitment processes, leading to high translational validity. However, the marked difference in maximal MEP magnitude emphasises the importance of species-specific adaptations.

## 1. Introduction

Cortical excitability (CE), defined as the strength of the response of cortical neurons to a given specific stimulation is meant to characterise the general reactivity of the neurons in a specific region (Ly et al., 2016). It is widely assessed by recording motor evoked potentials (MEP) in peripheral muscle fibres using electromyography (EMG) in response to single-pulse transcranial magnetic stimulation (sp-TMS), a method employed in both human studies and large-brain animal models, including non-human primates (NHPs).

More specifically, to quantify the responsiveness of the motor cortex from recorded MEPs, the motor threshold (MT) – traditionally defined as the minimum stimulation intensity to evoke a reliable motor response – has been introduced as a widely utilised measure in both basic research and clinical practice (Rossini et al., 1994, 2015). In practice, a “reliable motor response” is most often operationalized using an arbitrary criterion; by either 100 μV (Rossini et al., 1994; Tang et al., 2024) or 50 μV (Rossini et al., 2015; Ziemann et al., 2015) peak-to-peak MEP amplitude recorded over the target muscle. The most common MT determination methods include the relative frequency-based MT (fMT) approach, which is simple, intuitive, but is primarily aimed at distinguishing biological signals from the instrumental noise floor (Mills & Nithi, 1997; Wang et al., 2023), yielding a positively biased threshold value. More advanced probabilistic models (Awiszus, 2003; Wang et al., 2023), have also been developed for instance to account for the stochastic nature and non-Gaussian distribution of MEPs. However, most of these methods still estimate a one-dimensional index of CE and remain limited by the criterion-based conceptualization of a MT.

In line with this, studies have shown that traditionally measured MT values, regardless of the methodology utilised, are substantially higher than the minimum stimulation intensity required to evoke the first detectable MEP (Davies, 1984; Goetz et al., 2018; Li et al., 2022). Thus, while traditionally measured MT is valuable in establishing safety thresholds and rudimentarily personalising stimulation intensity ranges, advances in therapeutic techniques and neuroscience research necessitated MTs that are more physiologically grounded (Li et al., 2022). Recruitment curves (RCs), which describe the input-output properties of the corticospinal system (Devanne et al., 1997; Kukke et al., 2014) measure the relationship between stimulation intensity and MEP amplitude, thereby moving beyond the binary interpretation of a MEP response dictated by an arbitrary MT criterion. Beyond detecting small responses at low intensities (Davies, 1984; Goetz et al., 2018; Li et al., 2022), fitting a slope on the steepest part of the sigmoid curve, and considering its intercept with the floor of the sigmoid can yield a more principled description of excitability (Devanne et al., 1997; Kukke et al., 2014; Willer et al., 1987) . Hence, determining the stimulation intensity corresponding to this intercept where the RC turns from constant to logarithmic could define a physiologically relevant threshold.

Though recruitment curves and corticospinal excitability have been extensively researched in humans, preclinical research necessitates establishing translationally relevant animal models. Non-human primates (NHPs) are ideal candidates for such purpose due to their general similarity to humans, including direct cortico-motoneuronal projections to the hand and the similar anatomical organisation, comparable thickness (Lemon & Griffiths, 2005) and gyrification (Namba et al., 2019) of the neocortex with similar functional output. However, NHP studies remain limited in numbers (de Lima-Pardini et al., 2023), with most of them relying on the fMT methods based on visual observation of the muscle twitch or EMG recordings. Especially only a few studies have recorded RCs, and to the best of our knowledge those studies have been fully or partially conducted under general anaesthesia, which is known to alter CE and may prevent assessing the plateau phase of RC (Hanlon et al., 2021; Tang et al., 2024).

Here, we aimed to advance the characterization of CE in a preclinical translational framework that is meant to align with human standards. Using single-pulse TMS in awake rhesus macaques, we first measured traditionally defined MT through the relative frequency method. Subsequently, we recorded RCs in separate sessions and fitted Boltzmann sigmoid curves to allow for a more physiologically relevant and reproducible characterization of cortical excitability beyond the criterion-bound threshold method and provide a robust link between preclinical NHP animal models and human basic and clinical research.

## 2. Methods

### 2.1. Animals and housing

This study was performed using altogether 8 adult male rhesus macaque monkeys (*Macaca mulatta*), aged 12.4 (SD: 2.1) years (ranging from 9.1 to 16.9 years), weighing 8.8 (SD: 0.5) kg (ranging from 8.0 to 9.5 kg).

Animals were housed in pairs in home cages that were uniformly sized 230 × 100 × 200 cm (height × width × depth) and were equipped with wooden rest areas. In the vivarium and laboratories, temperature and humidity were maintained at 24 ± 1°C and 55 ± 5 RH% with continuous airflow. Light conditions were set to a 12/12 hour light-dark cycle (lights on at 07:00) using full spectrum lighting with transitions (twilight periods) between light and dark phases and were supplemented by natural illumination through side and roof windows to maintain the natural rhythms of the subjects. Following the daily training or testing sessions, subjects were fed once per day, in the afternoons. Diet was standard nutritionally complete laboratory chow specially designed for non-human primates (Altromin Spezialfutter GmbH) and was daily supplemented with fresh fruit and vegetables.

All procedures were conducted in the Grastyán Translational Research Centre of the University of Pécs. This study was approved by the Animal Welfare Committee of the University of Pécs and the Hungarian National Scientific Ethical Committee on Animal Experimentation. Ethical permission was issued by the Department of Animal Health and Food Control of the County Government Offices of the Ministry of Agriculture (BA02/2000-25/2020). Measures were taken to minimize pain and discomfort of the animals in accordance with the Directive 40/2013. (II.14.): ‘On animal experiments’ issued by the Government of Hungary, and the Directive 2010/63/EU ‘On the protection of animals used for scientific purposes’ issued by the European Parliament and the European Council.

### 2.2. Experimental set-up and initial training

A novel non-invasive head and arm fixation apparatus was developed as previously reported in Padányi et al. (2023) to facilitate frameless neuronavigation-assisted TMS stimulation. The apparatus consisted of an adjustable three-piece TMS face mask and an arm holder. The face mask stabilised the head while allowing access to the frontal, parietal and part of the occipital regions for TMS stimulation. Face masks were available in different sizes to accommodate the variation in the size of the subjects’ skull. The arm holder restricted the movement of the arms and the fingers separately, ensuring access to the abductor pollicis brevis muscle for electromyography (EMG) recording to enable the measurement of motor-evoked potentials (MEPs).

Following basic primate chair-training, all subjects were acclimatised to the laboratory environment, experimental conditions and set-up to facilitate awake, cooperative participation in experiments. Positive reinforcement training (PRT) with food rewards tailored to individual preferences was used. Full training and preparation (until motor threshold measurement) took 40.3 (SD: 13.9) sessions, out of which initial training and set-up habituation took 15.5 (SD: 4.6) sessions.

### 2.3. Transcranial magnetic stimulation (TMS) and electromyographic (EMG) recording

Focal magnetic stimulation was administered over the scalp using a MagPro x100 including MagOption (MagVenture, Lucernemarken, Denmark) and a figure-eight coil with active cooling (Cool-B65, MagVenture). The coil was manually positioned and kept tangentially over the hand area of the left motor cortex at an angle of 45 degrees to the sagittal plane, inducing a postero-anterior directional current flow.

Disposable surface electromyogram (EMG) electrodes (30 mm, Ag/AgCl, Covidien) were placed on the abductor pollicis brevis (APB) muscle of the right thumb in a tendon-belly montage, with the ground attached to the primate chair. Raw EMG waveform data was recorded in AC mode at 3 kHz sampling rate digitised at 12 bit with an analogue bandpass filter set between 16–470 Hz (EMG Pod, Rogue Research Inc., Canada). Data was saved in epochs between −50ms and +150ms relative to TMS pulse onset time (Brainsight™ TMS Frameless Navigation system, Rogue Research Inc, Montreal, Canada). MEP peak-to-peak amplitude was calculated in the 10 ms to 90 ms time window following stimulation in the BrainSight software.

### 2.4. Determination of primary motor hotspot

Prior to the start of the experiments each subject underwent a T1-weighted MRI scan to create the individual stereotaxic map of their brain. Brain images were imported into the Brainsight™ TMS Frameless Navigation system (Rogue Research Inc, Montreal, Canada) wherein head models and markers were created allowing for continuous online control of coil positioning during the experimental sessions using optical tracking of fiducial markers on the coil and on the face mask of the subject.

The APB muscle area was identified in the primary motor cortex (M1 hotspot) for each subject by systematic single-pulse biphasic stimulation over grid pattern centred around the expected M1 location on the precentral gyrus. The location yielding the largest and most consistent MEPs at a stable stimulation intensity (SI) was marked in the neuronavigation system and was further referred to as ‘M1 hotspot’. The M1 hotspot and the corresponding coil positioning target of an example subject is shown on **Figure 1**.

**Figure 1:**
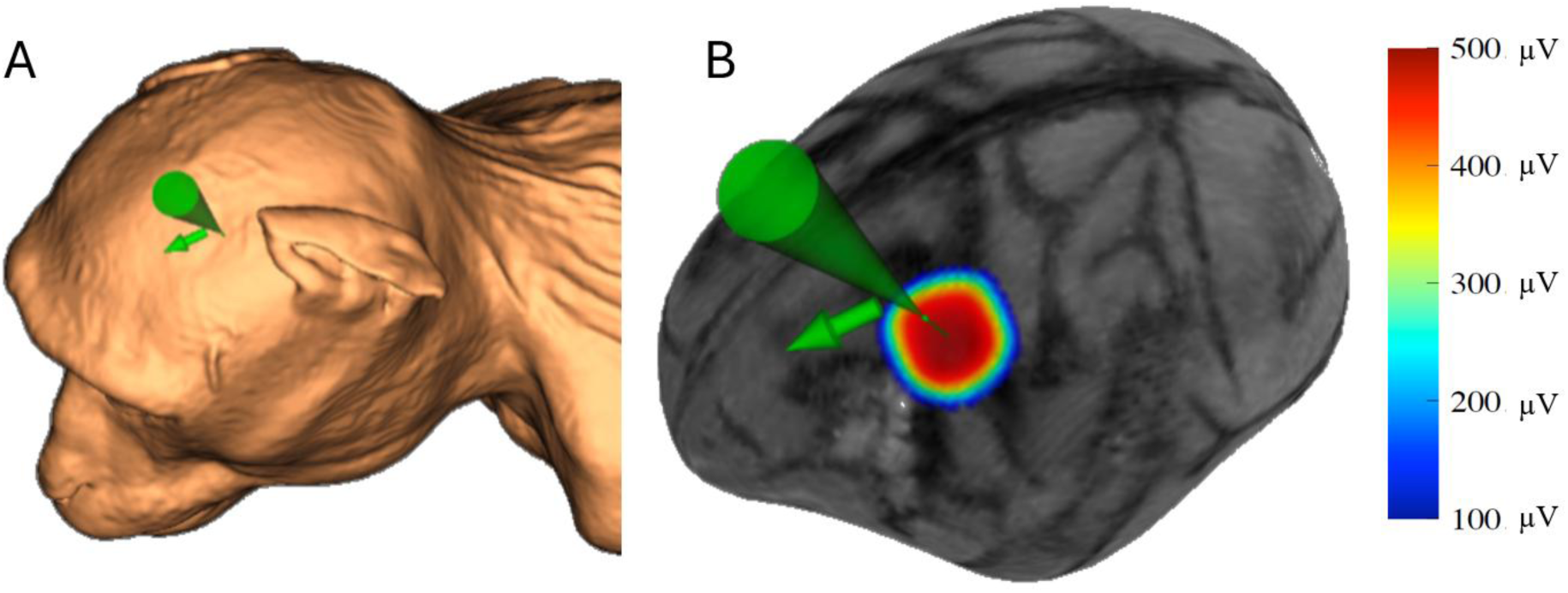
MRI-based reconstructions in the BrainSight neuronavigation software showing the position (green cone) and the orientation (green arrow) of the stimulation coil targeting the M1 hotspot in a single subject (‘Sp’). **A:** Skin reconstruction with coil position and orientation. **B:** Surface view of the cortex with overlayed heatmap as a smooth estimate of peak-to-peak MEP amplitudes recorded in a motor hotspot determination session. Right side: corresponding heatmap scale of MEP amplitudes.

At the beginning of the following motor threshold (MT) determination or recruitment curve (RC) recording sessions, the same M1 hotspot location was targeted, verified with a short, star-shaped stimulation pattern.

### 2.5. Determination of Motor Threshold (MT)

Traditional, criterion-based motor threshold (referred as tradMT) is generally defined as the minimum stimulation intensity at which the MEP peak-to-peak amplitude exceeds 50–100 µV in 50% of the trials (Rossini et al., 1994, 2015). In this study, tradMT was determined as the stimulation intensity at which at least 3 of 8 or 5 of 12 TMS pulses elicited MEPs meeting or exceeding the 100 µV criterion. Therefore, to find the MT at the marked M1 hotspot area, 8 to 12 pulses were delivered at a given simulator output intensity between 26% and 50% of maximum stimulator output (%MSO), using minimum 3 (maximum 9) stimulation levels in a staircase procedure which is in line with procedure describe by (Wang et al., 2023). The tradMT value for each subject (that was carried forward to the RC recording procedure described below) was calculated as the median of MT values obtained from all recorded and validated sessions. In one subject (‘Bat’) only 1 session was included, because in other sessions the protocol used did not reach high enough intensities to securely determine tradMT based on above given definition and stimulus intensity range. In addition, four sessions from three other subjects were discarded due to technical errors in EMG signal acquisition.

### 2.6. Recruitment curve recording

Recruitment curves (RCs), also known as input/output or stimulus/response curves, were recorded to characterise excitability of local motor cortical neurons and associated corticospinal tract fibres at nine relative stimulation intensities (relSI), expressed as multiples of the previously determined individual tradMT (0.5, 0.7, 0.8, 0.9, 1 = tradMT of the subject, 1.1, 1.2, 1.3, 1.5). Eight stimuli were delivered consecutively at the M1 hotspot for each stimulation intensity with at least 4-s inter-pulse interval. Stimulation intensities were administered in a semi-random order. We refer to the resulting dataset as ‘raw RC’, to differentiate from parametric recruitment curves, ‘fitted RCs’, explained below. Additionally, we measured MEP amplitudes at 1.2 × tradMT with 8 pulses right before and right after the RC recording.

### 2.7. Data analysis

Data analysis was conducted offline using R 4.3.2 (R Core Team, 2024), RStudio (RStudio Team, 2024). First, MEPs that were derived from pulses with a coil mispositioned or misaligned more than 1 mm or 3 degrees compared to the M1 hotspot target were deemed inappropriate and were excluded from further analysis. Then, root mean square (RMS) of baseline activity in each MEP epoch was calculated to quantify pre-pulse muscle activity. MEPs with a baseline RMS larger than 2 µV were deemed inappropriate and were excluded as the pulse was not delivered to the APB muscle at resting state.

Then, for the accepted MEP segments, peak-to-peak amplitudes were log transformed (with base 10). All analyses were conducted on log transformed MEPs. For visualisation, data and estimated parameters (RC MEP amplitude parameter estimates and endpoints of their confidence intervals) were back-transformed to linear µV values to be visualised on a logarithmic scale. Consequently, all averages/means on figures are geometric means (also equal to medians) on the original linear µV scale (Limpert et al., 2001), and MEP amplitude confidence intervals represent multiplicative uncertainty. Past work investigating the distribution of MEP amplitudes (Nielsen, 1996; Wassermann, 2002) and also advanced modelling studies (Goetz et al., 2014; Koponen et al., 2024) have shown that the log transformation partially mitigates the multiplicative variability of MEP data and is more suitable for our modelling procedure described below.

Using MEPs arising from the 1.2 × tradMT pulses administered before, during and after the RC, we probed for the possible cumulative excitability-changing effects of the single pulses delivered as part of the recruitment curve measurement. We used a linear mixed model with a factor of *Time* (3 levels: before, during, after RC) and random slopes for *Time* across sessions nested within subjects to check for an excitability shift. No significant excitability shift was found (F_2,3.9_=0.86, p=0.49).

A Boltzmann sigmoid function was fitted on the log-transformed peak-to-peak MEP amplitudes of each recorded RC separately, referred to as ‘fitted RC’. The Boltzmann sigmoid function (see e.g. Kukke et al, 2014) was defined as follows:

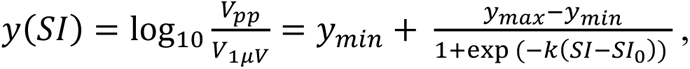

where *y(SI)* is log peak-to-peak amplitude as a function of stimulation intensity, *y_max_* is log maximum EMG amplitude (later referred to as ‘plateau of RC’), y_min_ is log minimum EMG amplitude, *SI_0_* is the inflection point, k is the slope and *V_1µV_* is the unit voltage (1 µV). The curve fitting was done in *R* using *nlminb* with the Hooke-Jeeves algorithm as a fallback method (*dfoptim* and *optimx* R packages). Note that due to the log transformation, the ‘linear’ phase of the sigmoid curve corresponds to an exponential gain on the original µV microvolt scale. In addition to the parameters of the Boltzmann curve, further points of interest were derived from the fit. We estimated the boundary points between the lower and higher asymptotic phases and the recruitment phase of the sigmoid curve, referred to as the lower and the higher “ankle points” of the sigmoid curve (*SI_low_* = *SI_0_*+2/*k*, and *SI_high_* = *SI_0_*−2/*k*, respectively). These ankle points were defined as stimulation intensities corresponding to the intersection of the asymptotes with the tangent line of the recruitment curve at the inflection point. The log MEP amplitudes corresponding to the ankle points were also calculated as the value of the sigmoid function at these points. Since lower ankle point represents the start of the recruitment phase of the curve, it can be used as a physiologically grounded estimate for the motor threshold (Devanne et al., 1997; Willer et al., 1987). We also estimated the stimulation intensity (SI) where the Boltzmann sigmoid function took the value of 100 µV as an approximation of the tradMT based on the fitted RC, referred to as ‘SI at 100µV recruitment’ (*SI_100µV_*).

To quantify the reliability of the tradMT and the fitted RC, separate random intercepts linear mixed models were fitted to the tradMT, the inflection point, the lower and upper ankle points and the SI at 100 µV recruitment, and intraclass correlation was calculated using the *performance* R package (‘0.11.0’, Lüdecke et al., 2021). Bootstrap confidence intervals with 95% coverage were calculated with 99999 iterations. To further characterize variability, intra- and inter-subject standard deviation parameters were also extracted from these models.

Next, we fitted group-level multivariate hierarchical linear models on the fitted RCs and the measured tradMTs. Two models were required to tackle numerical degeneracy arising from linear and nonlinear dependence between the fitted RC parameters – most of the fixed-effects parameters could be derived as linear contrasts of a smaller parameter set, but for hierarchical estimates and their variance-covariance parameters this would not be possible. The models were fitted using *lme4* (Bates et al., 2015) as nested hierarchical linear mixed models incorporating the variance-covariance structure among individual fitted RCs (or sessions) and animals (Dworkin & Bolker, 2023). First, a joint model was fitted that encompassed the whole four-parameter space of a fitted RC. For this, the session-level fitted RCs were reparametrized with two stimulation intensity parameters – the ankle-point SI-s, *SI_low_* and *SI_high_* – and two log MEP amplitude parameters – *y_max_* and *y_min_*. We also added tradMT to this model to be able to compare to fitted RC parameters. Fixed-effect parameters of interest were re-derived from the above joint group-level model as linear combinations of the included parameters, except from the *SI_100µV_* parameter (that was nonlinearly dependent). For estimating group-level hierarchical estimates and contrasts of *SI_100µV_*, an additional model of solely SI parameters were used with *SI_low_, SI_high_, SI_100µV_* and tradMT. Additionally, a variant of the joint model using *relSI* (×tradMT) instead of *SI* (%MSO) was fitted, since *relSI* is more comparable across subjects and also possibly across stimulation settings and hardware. For contrast estimates, degrees of freedom were approximated using Satterthwaite’s method.

All tests are two-tailed with a significance threshold of α=0.05, confidence intervals are with 95% coverage.

## 3. Results

The resting motor threshold measured in eight awake adult male rhesus macaques across 4.25±0.41 (mean±s.e.m.) sessions using the traditional method (Rossini et al., 2015) with 100 µV criterion (see **Methods**) was found to be reliable and stable (**Figure 2**). Within subject variability was small (SD: 2.41 %MSO) with an Intraclass Correlation (ICC) of 0.821 (95% CI: [0.44, 0.93]), which is considered to be solid (Kanig et al., 2023). As expected, variability between subjects was greater (SD: 5.16 %MSO) due to individual differences, with traditional MT (‘tradMT’) values ranging from 28% MSO to 46% MSO.

**Figure 2:**
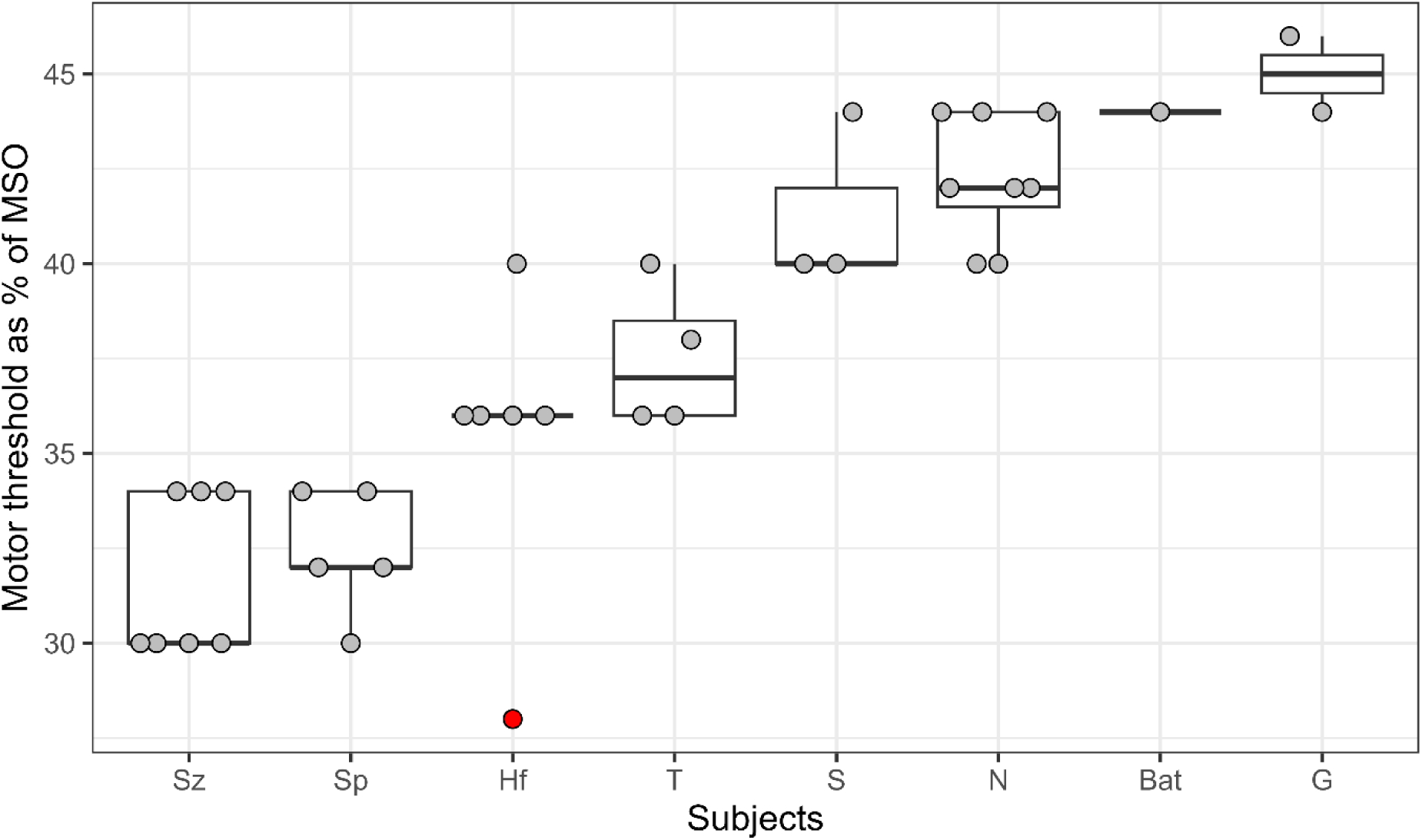
Traditional Motor Threshold (tradMT) and its variability in awake rhesus macaques (n=8). Boxplot of traditional motor threshold (displayed in percentage of Maximum Stimulation Output (%MSO) for each subject (n=8) showing median (horizonal line), interquartile range (box) and 1^st^ Quartile or 3^rd^ Quartile ± 1.5 × interquartile range (whiskers) where relevant. Single sessions are indicated by grey circles. Red indicates outlier, meaning outwith of the 1.5 × standard deviation (note that this data point was not excluded from analyses).

To characterize cortical excitability more comprehensively, Recruitment curves (raw RC) were recorded at various stimulation intensities expressed as multiples of the previously determined tradMT (0.5, 0.7, 0.8, 0.9, 1, 1.1, 1.2, 1.3, 1.5 × tradMT) in four subjects (**Figure 3A**). RCs were measured five times per subject to assess inter-session variability. Inspecting raw RC MEPs confirmed that meaningful MEPs with smaller amplitude were detected at stimulation intensities lower than the tradMT measured according to our standard protocol (Rossini et al., 1994, 2015), (**Figure 3D, Supplementary Material 1**), therefore we analysed RCs with a curve fitting procedure Specifically, a Boltzmann sigmoid function was fitted to each raw RC (for a representative single session with individual EMG signals, see **Figure 3B**; for a representative RC fit for the same session, see **Figure 3C**), yielding altogether n=20 fitted RCs (**Figure 3**). Beyond the four main parameters of the curve (plateau of RC, floor of RC, inflection point of RC, slope of RC), key CE parameters were derived: the lower and higher ankle points of RC, and the “SI at 100µV recruitment”, denoted *SI_100µV_* (**Figure 3B**, see **Methods** for details). Using separate linear mixed models for each parameter, we showed that SI parameters of the RC were stable and replicable (ICC_inflection point of RC_: 0.91, 95% CI: [0.37, 0.98], ICC_lower ankle point_: 0.84, 95% CI: [0.21, 0.96], ICC_higher ankle point:_ 0.79, 95% CI: [0.16, 0.94], ICC_SI at 100µVrecruitment_: 0.89, 95% CI: [0.33, 0.97]).

**Figure 3.**
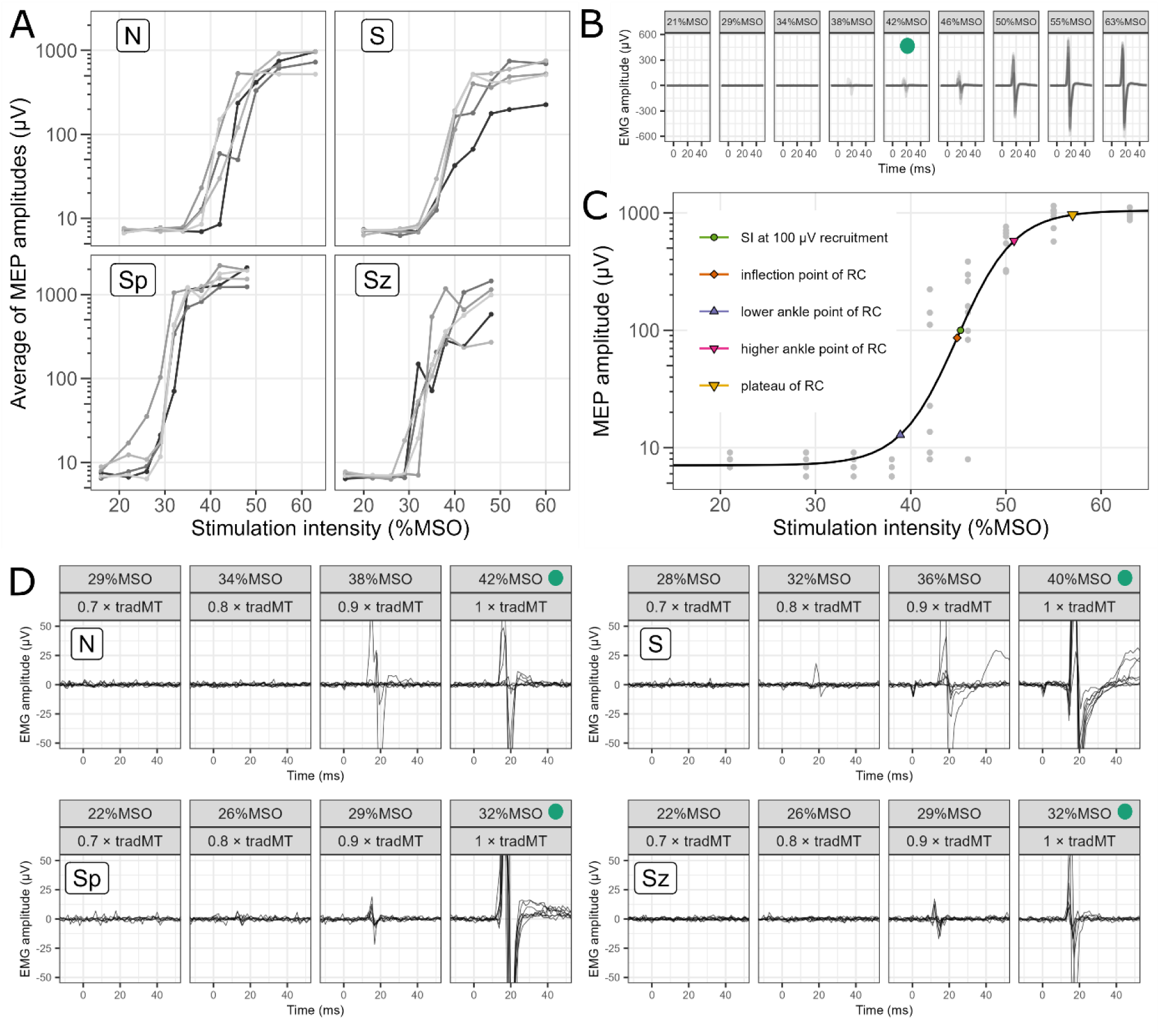
Individual MEPs and recruitment curves (RCs) recorded at multiple stimulation intensities (0.5 – 1.5 × tradMT) in n=4 subjects for n=5 sessions with example Boltzmann sigmoid curve fitting. **A:** Raw recruitment curves recorded at multiple stimulation intensities ranging from 0.5 × tradMT to 1.5 × tradMT for each subject. MEP amplitudes were averaged at each stimulation intensity level. The five sessions are displayed in different shades of grey. **B:** Motor evoked potentials in response to 8 pulses at each stimulation intensity (as %MSO) for one subject (‘N’) during an example session the with tradMT at 42 %MSO (marked by the filled green circle). **C:** Example of a fitted Boltzmann curve (a fitted RC) for subject (‘N’) during the same example session as in **B** with relevant RC parameters indicated with filled shapes. Grey points represent the MEP peak-to-peak amplitude of each stimulation. **D:** Individual MEPs of 8 pulses recorded at stimulation intensities 0.7, 0.8, 0.9 and 1 × tradMT (with %MSO values also shown) for all subjects during the sessions from the same day as example session in **B**. Note the presence of MEP responses below the tradMT, indicated by filled green circles.

Then we used a multivariate linear mixed model (mLMM, see **Methods**) to derive group-level estimates of the parameters and their comparisons against the tradMT. The group-level parameter estimates are visualised on **Figure 4B**, while individual conditional mean fitted RCs derived from the mLMM are plotted on **Figure 4A**. The tradMT (in the group mLMM: 36.5±3.0 %MSO) coincides well with both the inflection point (36.4±2.8 %MSO, comparison to tradMT: −0.12±0.9 %MSO, t_2.9_=−0.14, p=0.90, 95% CI: [−3.0, 2.7] %MSO) and its approximation based on the fitted RC, the SI_100µV_ (37.1±3.0 %MSO, comparison to tradMT: 0.6±0.8 %MSO, t_3.0_=0.79, p=0.49, CI: [−1.9, 3.1] %MSO), the latter of which provides an estimate for tradMT had it been measured in the session. Importantly, the lower ankle point was found to be on average at 32.7±2.4 %MSO, which was consistently and significantly below both the traditional MT (−3.8±1.1 %MSO, t_2.8_=−3.45, p=0.045, 95% CI: [−7.5, −0.15] %MSO) and SI_100µV_ (−4.4±0.9 %MSO, t_3.3_=−4.9, p=0.013, 95% CI: [−7.1, −1.7] %MSO).

**Figure 4.**
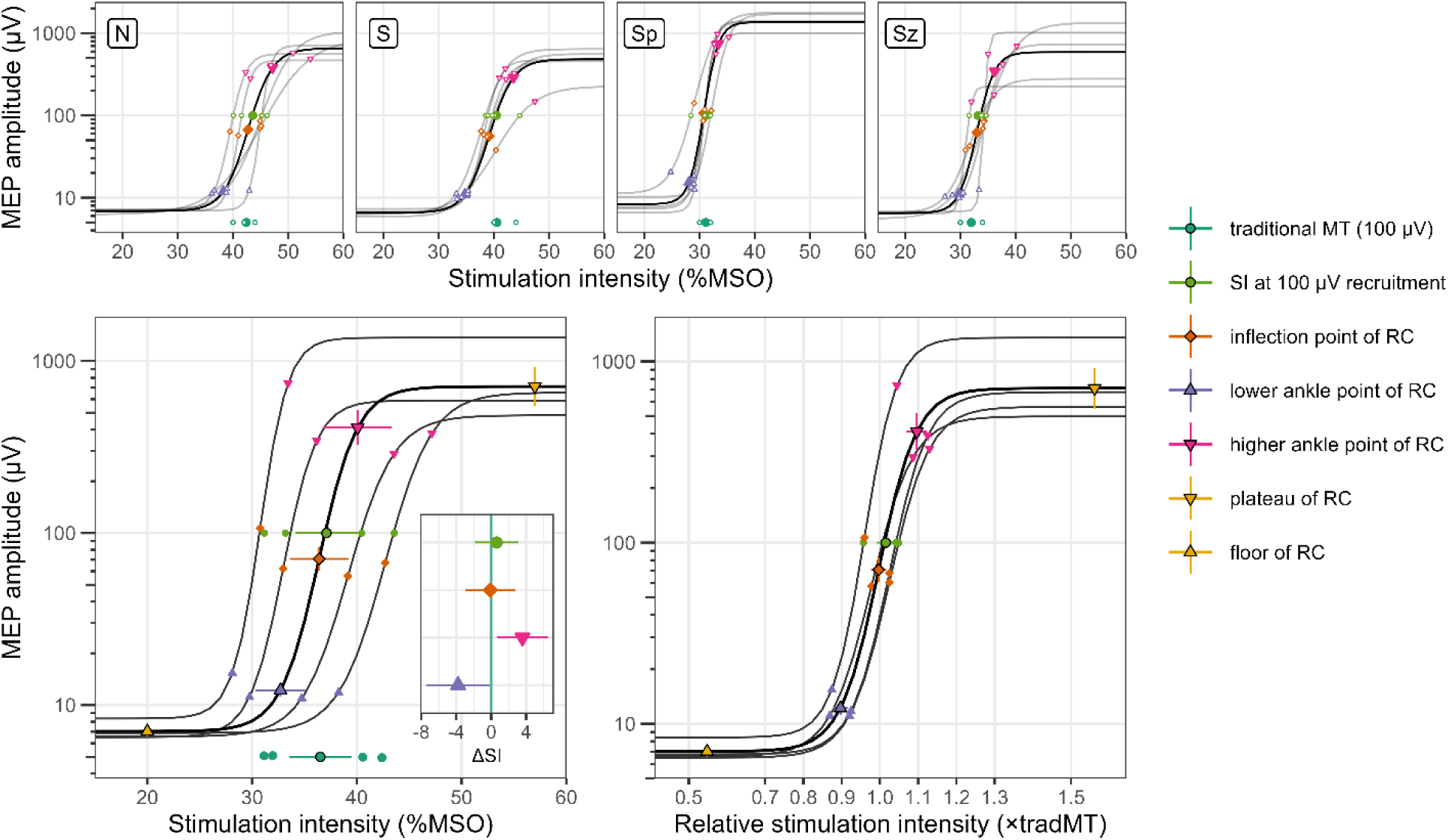
Fitted RCs and their parameters for individual sessions, subject averages and group average as a function of stimulation intensity (%MSO) and relative stimulation intensity (×tradMT). **A:** Fitted RCs for each subject. Individual session fitted RCs are shown as thin grey lines, bold black lines indicate subject averages. Fitted RC parameters of single sessions are shown with small, hollow shapes, while the large, filled shapes indicate subject average parameters. **B:** Subject average (thin grey lines) and grand average (bold black line) fitted RCs with parameters (filled shapes for subject averages, filled shapes with contour showing group average). The inset shows group-level contrast estimates with 95% CIs for the difference between tradMT (0 on x axis) and ankle points, inflection point and SI_100µV_.

Accordingly, the higher ankle point (40.1±3.3 %MSO) was consistently above the tradMT (3.6±1.1 %MSO, t_4.0_=3.40, p=0.027, 95% CI: [0.7, 6.5] %MSO).

Since stimulation intensities are frequently expressed relative to a traditionally determined MT, we also generated mLMM-based summaries for relative stimulation intensities. The lower ankle point was at 0.90±0.02 × tradMT (CI: [0.84, 0.96] × tradMT), and the higher ankle point was at 1.10±0.03×tradMT (CI: [1.02, 1.18] × tradMT).

In agreement with group-level results from the fitted RC, a comprehensive overview of all recorded MEPs and corresponding fitted RC parameters in **Supplementary Material 1** confirms that in the majority of individual sessions we measured well-formed MEPs around or above the lower ankle point, below S_I100µV_ (and thereby, also below the tradMT).

We also estimated group-level expectations of the most important MEP amplitude parameters, the fitted RC plateau and the lower ankle point (**Figure 4**, y coordinates of parameter points). The plateau of the fitted RC was on average at 711 µV (95% CI: [314, 1611] µV), ranging from 224 µV to 1790 µV. The lower ankle point was found at amplitudes ranging from 10 to 21 µV (hierarchical model estimate for group mean: 12.2 µV, 95% CI: [9.3, 16.0] µV), considerably lower than commonly observed criterion levels of 100 or 50 µV.

## 4. Discussion

We probed cortical excitability (CE) using single-pulse transcranial magnetic stimulation (sp-TMS) in awake, non-invasively restricted, fully cooperative rhesus macaques. Beyond measuring the traditionally defined motor threshold (referred to as ‘tradMT’), which was found to be stable and replicable, we recorded recruitment curves (raw RC) and fitted them with the Boltzmann sigmoid (fitted RC) function. The parameters of the fitted curve provided a more comprehensive description of motor cortical excitability compared to the one-dimensional tradMT measure. Notably, the tradMT values corresponded most closely with the inflection point of the curve, while physiologically relevant threshold values were identified based on the lower ankle point of the fitted curve.

To directly compare parameters of the fitted RC and MTs based on a 100 µV criterion, we measured tradMT values using the relative frequency method, and we also estimated the ‘SI at 100 µV recruitment’ (*SI_100µV_*) parameter based on the RC, which is more comparable to other advanced or model-driven methods to determine criterion-based MT (Awiszus, 2003; Mills & Nithi, 1997; Wang et al., 2023). Additionally, *SI_100µV_* did not differ from tradMT, confirming consistent cortical excitability across the disparate tradMT and RC measurement sessions. Consequently, comparison between parameters of the sigmoid function and the tradMT derived from separate sessions is justified and discussed further below.

Firstly, motor evoked potential (MEP) responses with the expected morphology occurred at stimulation intensities well below the tradMT values. They were much smaller in amplitude, in line with Hennemann’s size principle (1957), which accounts for the recruitment of smaller motor units at lower stimulation intensities. Small-amplitude near-threshold MEPs can be overlooked even if reliably measured and detected, due to the multiplicative scaling of MEPs and the ensuing non-Gaussian amplitude distribution. To obtain a physiological threshold based on the fitted RC, we determined the ‘lower ankle point of the RC’ as the point where the RC transitions from a constant floor to a logarithmic phase (that is apparently linear on our figures with log scale). This concept is analogous to thresholds described in previous studies using regression methods (Devanne et al., 1997; Goetz et al., 2018; Tyagi et al., 2024). The lower ankle point was very close to 0.9 × tradMT (also termed as 90% or 0.9 relative stimulation intensity, relSI) for all four animals, which is a substantial difference that amounts to half the dynamic range of the recruitment phase (with the higher ankle point found at 1.1 × tradMT). In turn, tradMT and *SI_100µV_* were found near the inflection point of the curve (on a log scale), consistently and significantly above the physiological threshold, as also reported for humans in recent studies (Li et al., 2022; Tyagi et al., 2024). While both the inflection point and the physiological threshold are valuable metrics for describing cortical excitability, they represent distinct characteristics of the recruitment process (see also Tyagi et al, 2024). Displaying and analysing RCs on a logarithmic scale highlights these distinctions. The inflection point of the RCs and the tradMT are located at the midpoint of the input-output dynamic range (represented linearly on a logarithmic scale), thus here the motor response amplitude (i.e., MEP) is the most sensitive to the changes in excitability, yet they can still be reliably inducible. Therefore, tradMT remains a robust CE index that is generalizable across stimulation protocols. At the same time, similarly to humans, we found that in rhesus macaques the tradMT also does not correspond to a physiological threshold. So, even newer approaches with advanced methods (Wang et al., 2023) might provide more accurate estimates, these advanced methods still remain tied to arbitrary criterion-based thresholds established exclusively in humans. TradMT and similar CE indices also do not extend beyond the one-dimensional measure associated with the inflection point of the RC, and fail to provide additional insights into recruitment dynamics.

Importantly, the shape and the characteristics of the RCs are remarkably similar in humans and rhesus macaques suggesting a shared underlying recruitment process. However, the present results highlight and quantify important differences between macaques and humans that are of key importance with respect to translatability. Firstly, in the present study, the tradMT values in awake rhesus macaques occurred at 37.5±0.9% of maximum stimulator output (MSO) with the inflection point of the RCs as 36.3±2.8 %MSO. By contrast, with the same equipment as in our study (C-B65 coil and a MagVenture stimulator), Hanlon et al. (2021) reported higher values (76.7 %MSO ± 5.1% SD) at the left hand in female rhesus macaques under anesthesia, reaffirming the importance of cortical state for excitability measurement. Direct comparison with human studies is hindered by the variety of stimulators and coils used, underscoring the need for more wide-spread utilization of e-field modelling in both humans and NHPs (Dannhauer et al., 2024; Thielscher et al., 2015). Nonetheless, in the closest possible up-to-date contrast Jitsakulchaidej et al. (2022) reported 47.31 ± 8.06% SD in the APB muscle (MMC-140-II parabolic coil and MagVenture stimulator).

Another important result with respect to translational TMS applications is the plateu RC amplitude. Peak MEP amplitudes (the RC plateau) in awake male rhesus macaques in our study ranged from approx. 700 to 1200 µV, substantially higher than 300-400 µV of peak amplitude measured for male rhesus macaques under anaesthesia (Tang et al., 2024) or the 180-480 µV peak-to-peak amplitude for female rhesus macaques under anaesthesia (Hanlon et al., 2021). It also presents a stark contrast to human results, which report even higher plateaus, for instance the 2.5 mV in Rosenkranz et al. (2007) for the APB muscle or 4.5 mV in Kukke et al. (2014) for FDI muscle or the 10 mV in Koponen et al. (2024) similarly in the FDI muscle. These differences may be attributed to the greater fascicle length, and denser muscle innervation in humans compared to macaques (Paul, 2001).

Thus, while the recruitment processes appear fundamentally similar between species, it is crucial to take note of the magnitude differences, especially when establishing one-dimensional criteria for stimulation protocols. Such is the case for paired-pulse protocols and subsequent RC recording, where for humans the range of stimulation intensities are centred around the intensity where 1 mV MEP occurred. Our results are also relevant for setting stimulation intensity in repeated TMS and theta burst protocols (Tyagi et al., 2024).

A key strength of our study was the training and experimental set-up which enabled RC recording (beyond simple criterion-based MT protocols) in awake rhesus macaques with minimal discomfort. One might argue that due to the lack of anaesthesia, the varying muscle tension or tone between stimulation pulses and/or sessions could be a potential confounding factor. To address this, we monitored baseline muscle activity and excluded segments exceeding the 2 µV RMS threshold (see Methods) to ensure that the RC recordings were not contaminated or confounded by extraneous muscle activity.

In conclusion, we demonstrated the feasibility of recording stable and replicable tradMT and RC measures in awake rhesus macaques. Furthermore, parameters derived from Boltzmann sigmoidal curve fitted on the raw RCs revealed distinction between tradMT, which aligns with the inflection point of the curve, and the physiological threshold at the lower ankle point of the curve. While the overall RC shape was consistent with human data suggesting that recruitment processes are underlined by the same or very similar mechanisms leading to high translational validity. Moreover, the observed difference in magnitude emphasises the importance of species-specific adaptations, which is pertinent not only to basic research but also to preclinical and clinical applications across various fields, including neurodegenerative and psychiatric disorders.

## Acknowledgements

The authors would like to thank Szuhád Khalil and Evelin Kiefer, for their valuable technical contribution during equipment preparation and data acquisition.

## Author contributions

A.P.: Conceptualization, Methodology, Investigation, Data Curation, Formal analysis, Visualization, Writing – Original Draft, Writing – Review and Editing;

B.K.: Conceptualization, Methodology, Software, Data Curation, Formal analysis, Visualization, Writing – Original Draft, Writing – Review and Editing;

B.L.: Resources, Project administration, Funding acquisition, Writing – Review and Editing;

I.H.: Conceptualization, Methodology, Resources, Writing – Original Draft, Writing – Review and Editing, Supervision, Project administration, Funding acquisition.

## References

Awiszus, F. (2003). Chapter 2 TMS and threshold hunting. In Supplements to Clinical Neurophysiology (Vol. 56, pp. 13–23). Elsevier. 10.1016/S1567-424X(09)70205-3

Bates, D., Mächler, M., Bolker, B., & Walker, S. (2015). Fitting Linear Mixed-Effects Models Using lme4 | Journal of Statistical Software. *Journal of Statistical Software*, *Vol 65*(67(1)), 1–48. 10.18637/jss.v067.i01

Dannhauer, M., Gomez, L. J., Robins, P. L., Wang, D., Hasan, N. I., Thielscher, A., Siebner, H. R., Fan, Y., & Deng, Z.-D. (2024). Electric Field Modeling in Personalizing Transcranial Magnetic Stimulation Interventions. Biological Psychiatry, 95(6), 494–501. 10.1016/j.biopsych.2023.11.022

Davies, T. W. (1984). Definition of human reflex excitability by statistical analysis of quantal EMG responses. Brain Research, 293(2), 386–389. 10.1016/0006-8993(84)91249-6

de Lima-Pardini, A. C., Mikhail, Y., Dominguez-Vargas, A.-U., Dancause, N., & Scott, S. H. (2023). Transcranial magnetic stimulation in non-human primates: A systematic review. Neuroscience & Biobehavioral Reviews, 152, 105273. 10.1016/j.neubiorev.2023.105273

Devanne, H., Lavoie, B. A., & Capaday, C. (1997). Input-output properties and gain changes in the human corticospinal pathway. Experimental Brain Research, 114(2), 329–338. 10.1007/PL00005641

Dworkin, I., & Bolker, B. (2023, November 6). Multivariate modeling via mixed models. https://web.archive.org/web/20241125140301/https://mac-theobio.github.io/QMEE/lectures/MultivariateMixed.notes.html

Goetz, S. M., Li, Z., & Peterchev, A. V. (2018). Noninvasive Detection of Motor-Evoked Potentials in Response to Brain Stimulation Below the Noise Floor—How Weak Can a Stimulus Be and Still Stimulate. 2018 40th Annual International Conference of the IEEE Engineering in Medicine and Biology Society (EMBC), 2687–2690. 10.1109/EMBC.2018.8512765

Goetz, S. M., Luber, B., Lisanby, S. H., & Peterchev, A. V. (2014). A Novel Model Incorporating Two Variability Sources for Describing Motor Evoked Potentials. Brain Stimulation, 7(4), 541–552. 10.1016/j.brs.2014.03.002

Hanlon, C. A., Czoty, P. W., Smith, H. R., Epperly, P. M., & Galbo, L. K. (2021). Cortical excitability in a nonhuman primate model of TMS. *Brain Stimulation: Basic*, Translational, and Clinical Research in Neuromodulation, 14(1), 19–21. 10.1016/j.brs.2020.10.008

Henneman, E. (1957). Relation between Size of Neurons and Their Susceptibility to Discharge. Science. 10.1126/science.126.3287.1345

Jitsakulchaidej, P., Wivatvongvana, P., & Kitisak, K. (2022). Normal parameters for diagnostic transcranial magnetic stimulation using a parabolic coil with biphasic pulse stimulation. BMC Neurology, 22, 510. 10.1186/s12883-022-02977-8

Kanig, C., Osnabruegge, M., Schwitzgebel, F., Litschel, K., Seiberl, W., Mack, W., Schoisswohl, S., & Schecklmann, M. (2023). Retest reliability of repetitive transcranial magnetic stimulation over the healthy human motor cortex: A systematic review and meta-analysis. Frontiers in Human Neuroscience, 17. https://www.frontiersin.org/articles/10.3389/fnhum.2023.1237713

Koponen, L. M., Martinez, M., Wood, E., Murphy, D. L. K., Goetz, S. M., Appelbaum, L. G., & Peterchev, A. V. (2024). Transcranial magnetic stimulation input–output curve slope differences suggest variation in recruitment across muscle representations in primary motor cortex. Frontiers in Human Neuroscience, 18, 1310320. 10.3389/fnhum.2024.1310320

Kukke, S. N., Paine, R. W., Chao, C., de Campos, A. C., & Hallett, M. (2014). Efficient and reliable characterization of the corticospinal system using transcranial magnetic stimulation. Journal of Clinical Neurophysiology : Official Publication of the American Electroencephalographic Society, 31(3), 246–252. 10.1097/WNP.0000000000000057

Lemon, R. N., & Griffiths, J. (2005). Comparing the function of the corticospinal system in different species: Organizational differences for motor specialization? Muscle & Nerve, 32(3), 261–279. 10.1002/mus.20333

Li, Z., Peterchev, A. V., Rothwell, J. C., & Goetz, S. M. (2022). Detection of Motor-Evoked Potentials Below the Noise Floor: Rethinking the Motor Stimulation Threshold. Journal of Neural Engineering, 19(5), 10.1088/1741-2552/ac7dfc. https://doi.org/10.1088/1741-2552/ac7dfc

Limpert, E., Stahel, W. A., & Abbt, M. (2001). Log-normal Distributions across the Sciences: Keys and Clues: On the charms of statistics, and how mechanical models resembling gambling machines offer a link to a handy way to characterize log-normal distributions, which can provide deeper insight into variability and probability—normal or log-normal: That is the question. BioScience, 51(5), 341–352. 10.1641/0006-3568(2001)051[0341:LNDATS]2.0.CO;2

Ly, J. Q. M., Gaggioni, G., Chellappa, S. L., Papachilleos, S., Brzozowski, A., Borsu, C., Rosanova, M., Sarasso, S., Middleton, B., Luxen, A., Archer, S. N., Phillips, C., Dijk, D.-J., Maquet, P., Massimini, M., & Vandewalle, G. (2016). Circadian regulation of human cortical excitability. Nature Communications, 7(1), 11828. 10.1038/ncomms11828

Mills, K. R., & Nithi, K. A. (1997). Corticomotor threshold to magnetic stimulation: Normal values and repeatability. Muscle & Nerve, 20(5), 570–576. 10.1002/(sici)1097-4598(199705)20:5<570::aid-mus5>3.0.co;2-6

Namba, T., Vaid, S., & Huttner, W. B. (2019). Primate neocortex development and evolution: Conserved versus evolved folding. Journal of Comparative Neurology, 527(10), 1621–1632. 10.1002/cne.24606

Nielsen, J. F. (1996). Logarithmic Distribution of Amplitudes of Compound Muscle Action Potentials Evoked by Transcranial Magnetic Stimulation. Journal of Clinical Neurophysiology, 13(5), 423.

Padányi, A., Knakker, B., Kiefer, E., Khalil, S., & Hernádi, I. (2023, August 28). Investigation of cortical excitability with non-invasive transcranial magnetic stimulation in awake non-human primates. European Brain and Behaviour Society Conference, Amsterdam.

Paul, A. C. (2001). Muscle length affects the architecture and pattern of innervation differently in leg muscles of mouse, guinea pig, and rabbit compared to those of human and monkey muscles. The Anatomical Record, 262(3), 301–309. 10.1002/1097-0185(20010301)262:3<301::AID-AR1045>3.0.CO;2-H

Rosenkranz, K., Kacar, A., & Rothwell, J. C. (2007). Differential Modulation of Motor Cortical Plasticity and Excitability in Early and Late Phases of Human Motor Learning. The Journal of Neuroscience, 27(44), 12058. 10.1523/JNEUROSCI.2663-07.2007

Rossini, P. M., Barker, A. T., Berardelli, A., Caramia, M. D., Caruso, G., Cracco, R. Q., Dimitrijević, M. R., Hallett, M., Katayama, Y., & Lücking, C. H. (1994). Non-invasive electrical and magnetic stimulation of the brain, spinal cord and roots: Basic principles and procedures for routine clinical application. Report of an IFCN committee. Electroencephalography and Clinical Neurophysiology, 91(2), 79–92. 10.1016/0013-4694(94)90029-9

Rossini, P. M., Burke, D., Chen, R., Cohen, L. G., Daskalakis, Z., Di Iorio, R., Di Lazzaro, V., Ferreri, F., Fitzgerald, P. B., George, M. S., Hallett, M., Lefaucheur, J. P., Langguth, B., Matsumoto, H., Miniussi, C., Nitsche, M. A., Pascual-Leone, A., Paulus, W., Rossi, S., … Ziemann, U. (2015). Non-invasive electrical and magnetic stimulation of the brain, spinal cord, roots and peripheral nerves: Basic principles and procedures for routine clinical and research application. An updated report from an I.F.C.N. Committee. Clinical Neurophysiology, 126(6), 1071–1107. 10.1016/j.clinph.2015.02.001

Tang, C., Zhang, W., Zhang, X., Zhou, J., Wang, Z., Zhang, X., Wu, X., Su, H., Jiang, H., Zhai, R., & Zhao, M. (2024). A 3D-Printed helmet for precise and repeatable neuromodulation targeting in awake non-human primates. Heliyon, 10(17), e37121. 10.1016/j.heliyon.2024.e37121

Thielscher, A., Antunes, A., & Saturnino, G. B. (2015). Field modeling for transcranial magnetic stimulation: A useful tool to understand the physiological effects of TMS? 2015 37t*h Annual International Conference of the IEEE Engineering in Medicine and Biology Society (EMBC)*, 222–225. 10.1109/EMBC.2015.7318340

Tyagi, V., Murray, L. M., Asan, A. S., Mandigo, C., Virk, M. S., Harel, N. Y., Carmel, J. B., & McIntosh, J. R. (2024). *Hierarchical Bayesian estimation of motor-evoked potential recruitment curves yields accurate and robust estimates* (No. arXiv:2407.08709). arXiv. 10.48550/arXiv.2407.08709

Wang, B., Peterchev, A. V., & Goetz, S. M. (2023). Three novel methods for determining motor threshold with transcranial magnetic stimulation outperform conventional procedures. Journal of Neural Engineering, 20(5), 056002. 10.1088/1741-2552/acf1cc

Wassermann, E. M. (2002). Variation in the response to transcranial magnetic brain stimulation in the general population. Clinical Neurophysiology, 113(7), 1165–1171. 10.1016/S1388-2457(02)00144-X

Willer, J. C., Miserocchi, G., & Gautier, H. (1987). Hypoxia and monosynaptic reflexes in humans. Journal of Applied Physiology, 63(2), 639–645. 10.1152/jappl.1987.63.2.639

Ziemann, U., Reis, J., Schwenkreis, P., Rosanova, M., Strafella, A., Badawy, R., & Müller-Dahlhaus, F. (2015). TMS and drugs revisited 2014. Clinical Neurophysiology, 126(10), 1847–1868. 10.1016/j.clinph.2014.08.028

