## Supplementary Material 1 for "Assessment of cortical excitability in awake rhesus macaques with transcranial magnetic stimulation: translational insights from recruitment curves"

*for*

Version: 2024.12.11

This supplementary document contains individual Motor Evoked Potentials (MEPs) for each raw recruitment curve (RC) of each animal, focusing on near-threshold, small-amplitude responses. Each figure represents the recruitment curve (RC) recording from a given session of one animal. At the bottom, a collective x axis shows the TMS Stimulation Intensity (SI) in % of maximal stimulator output (%MSO) and as a multiple of the traditional motor threshold (×tradMT). Above, 9 columns of MEP traces are laid out positioned along the x axis according to the Stimulation Intensity applied, with 8-10 MEP traces in each column (16 in one case). To emphasise small amplitude responses, MEP trace y axes are clipped at ±20 µV; the time axis spans [-10, 50] ms – scale bars for both axes are found at the bottom left. A black arrow marks t=0 ms, the time point of stimulation.

Just above the collective x axis the tradMT and 4 fitted RC parameters are shown as coloured markers; the corresponding legend at the bottom also shows SI values and displayed in order of increasing SI to facilitate comparison. TradMTs were measured in separate sessions preceding RC recording, but importantly, the SI at 100 µV recruitment ($SI_{100\mu V}$) parameter of the fitted RC (indicated as SI at 100µV on the figure) provides an estimate for what the tradMT would be if it was measured in the same session. The results of the group analysis of the main text can be confirmed in each session: the lower ankle point (a physiologically grounded motor threshold) is lower than the $SI_{100\mu V}$ (and thereby, the tradMT), and there are well-formed MEPs around or above the lower ankle point, below the 100 µV criterion level.

### Animal 'N', Session 1

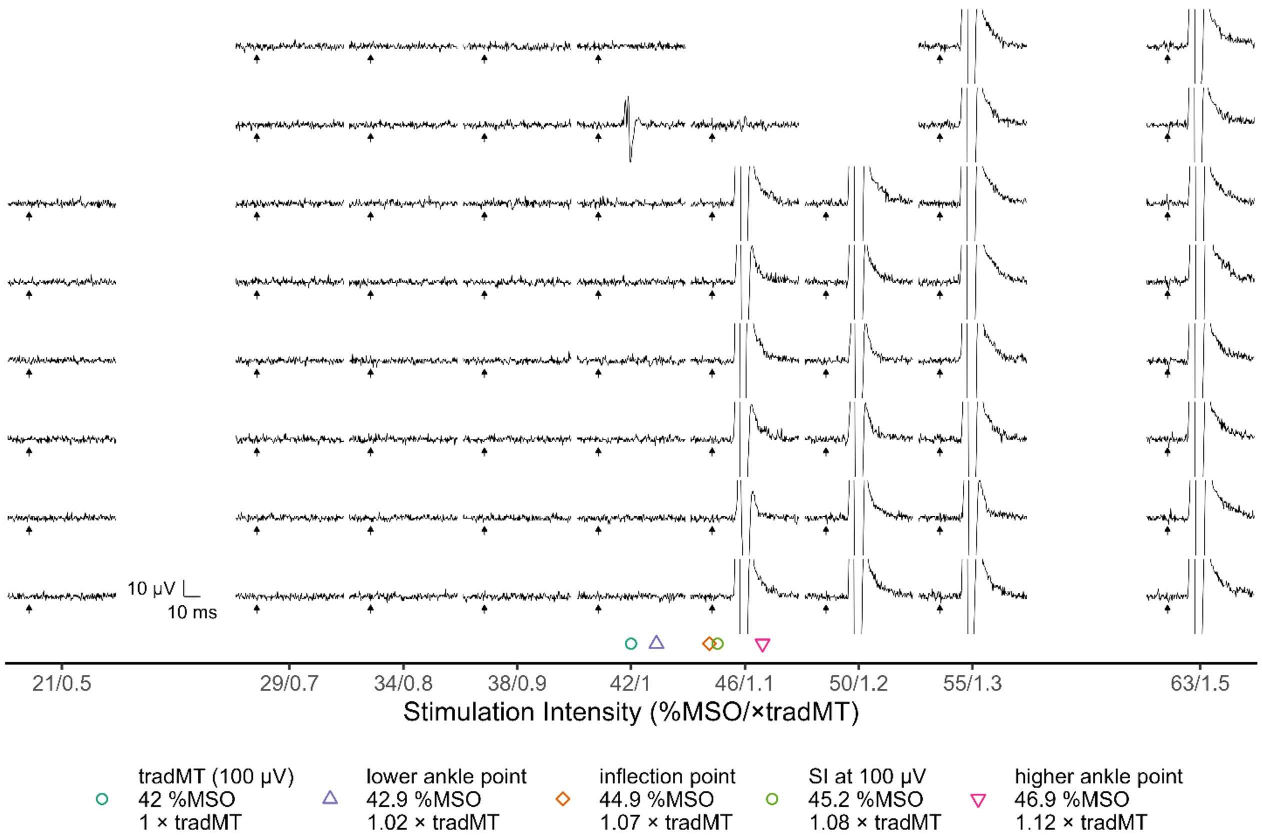


### Animal 'N', Session 2

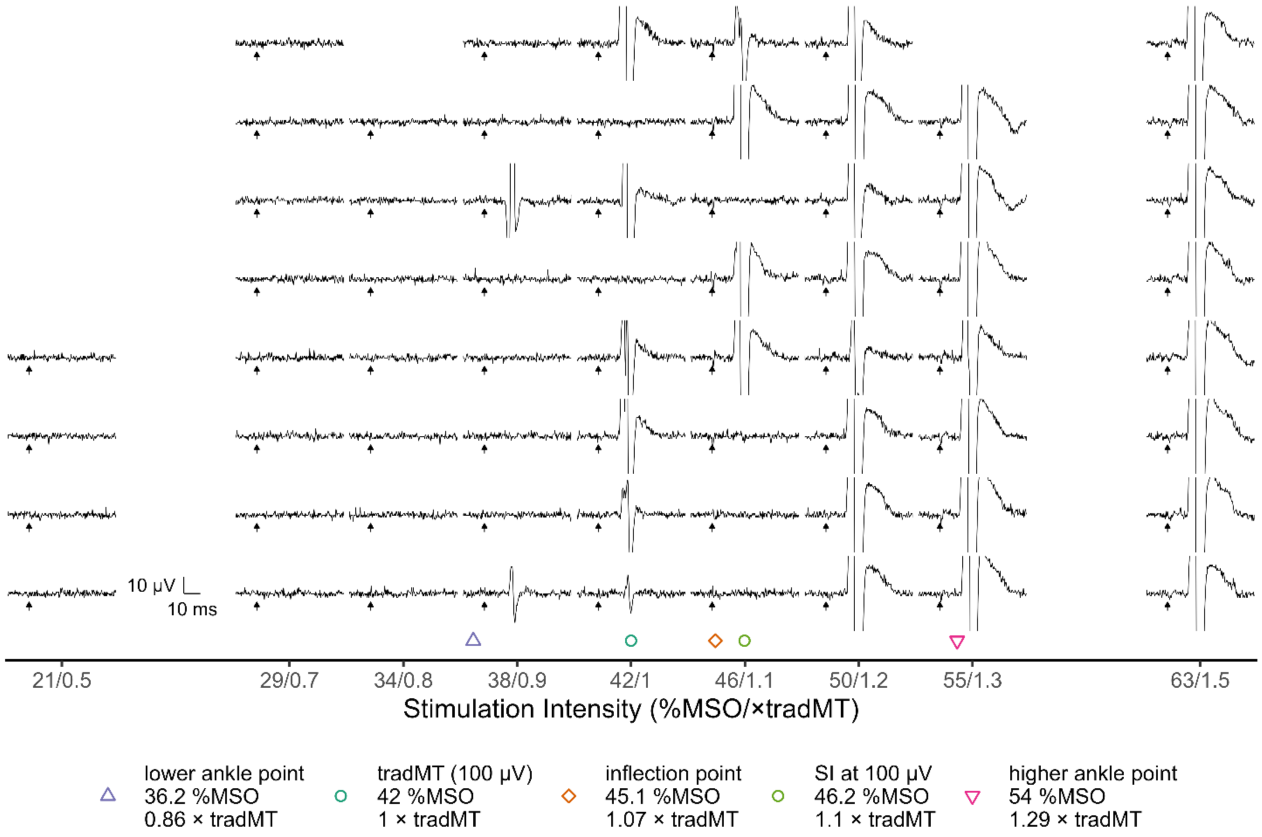

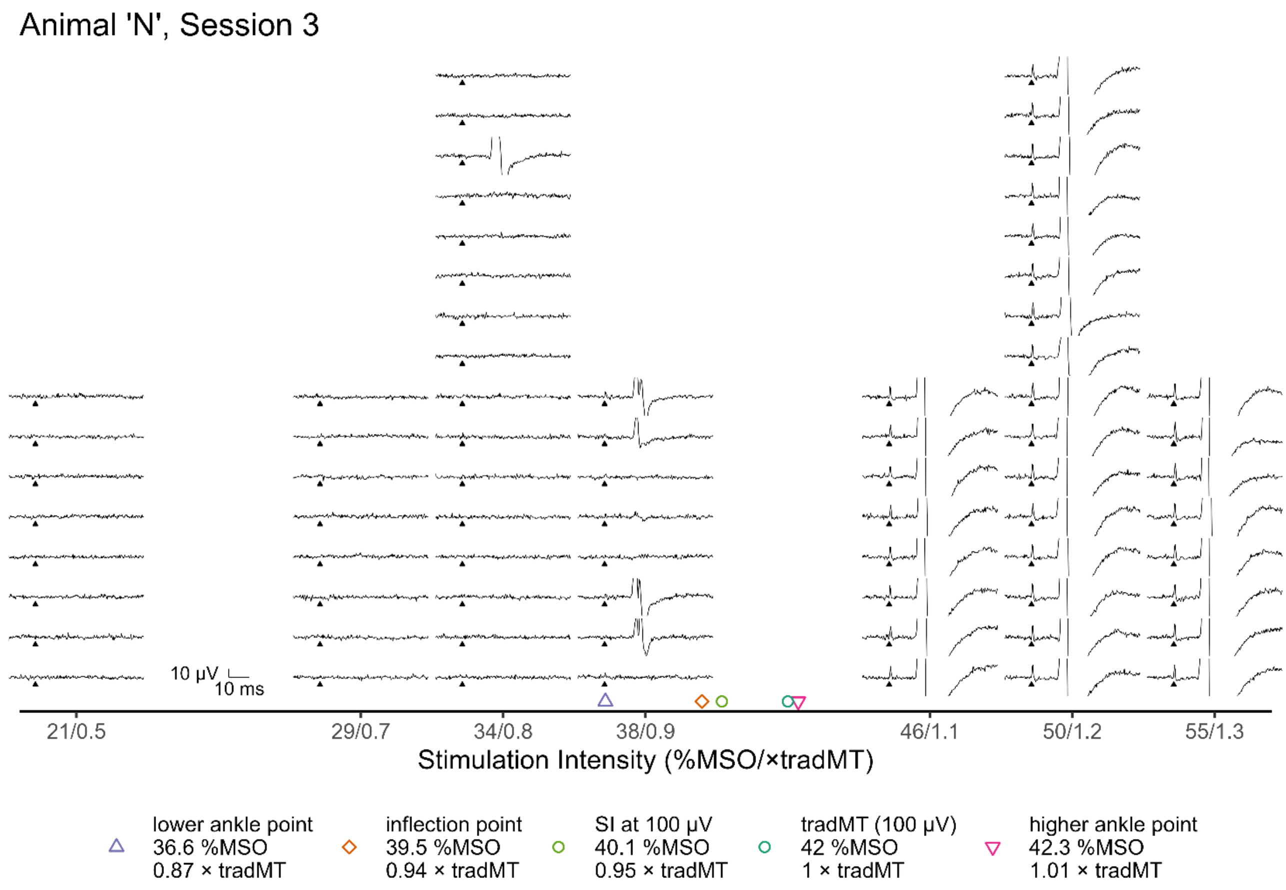
Animal 'N', Session 3
10 µV
10 ms
21/0.5
29/0.7
34/0.8
38/0.9
46/1.1
50/1.2
55/1.3
Stimulation Intensity (%MSO/×tradMT)
lower ankle point
36.6 %MSO
0.87 × tradMT
inflection point
39.5 %MSO
0.94 × tradMT
SI at 100 µV
40.1 %MSO
0.95 × tradMT
tradMT (100 µV)
42 %MSO
1 × tradMT
higher ankle point
42.3 %MSO
1.01 × tradMT

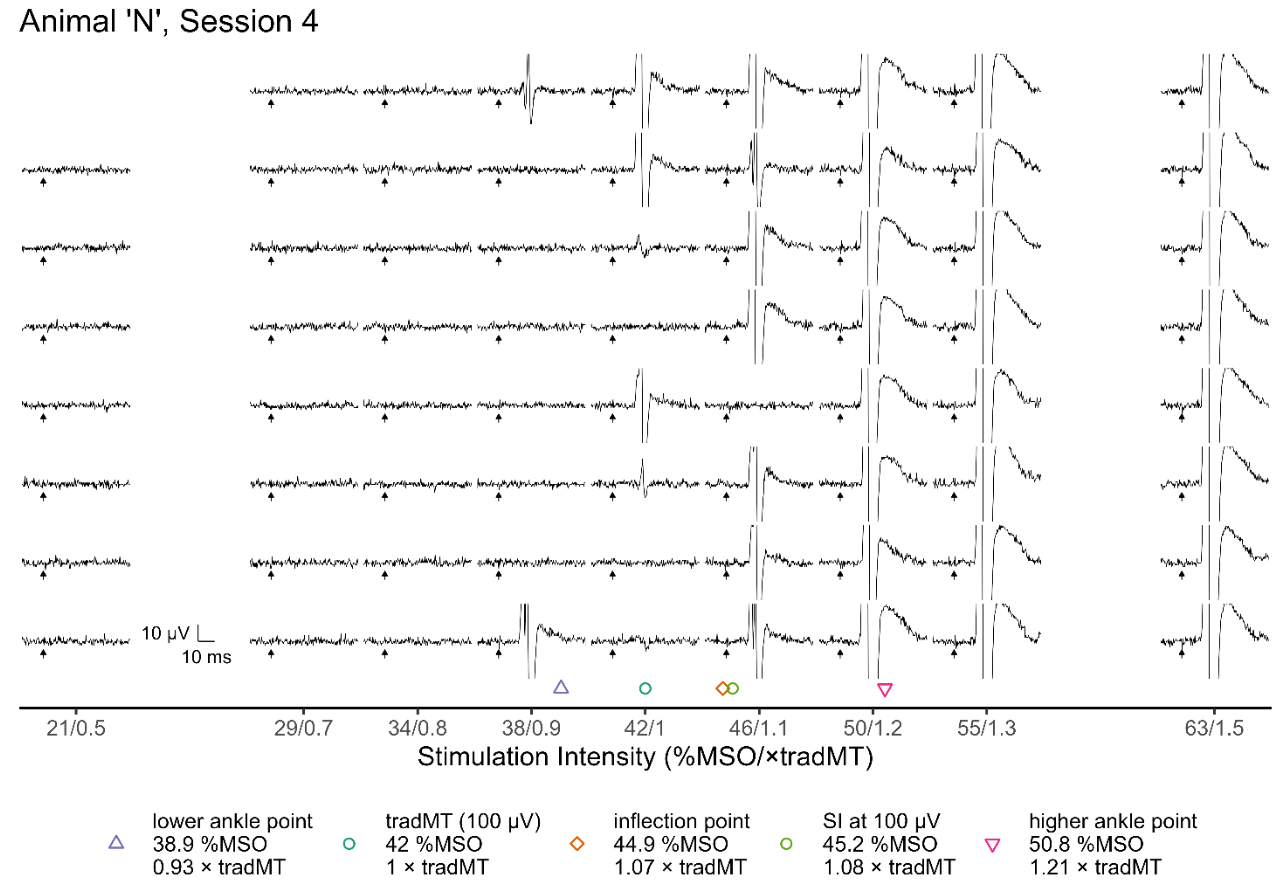
Animal 'N', Session 4
10 µV
10 ms
21/0.5
29/0.7
34/0.8
38/0.9
42/1
46/1.1
50/1.2
55/1.3
63/1.5
Stimulation Intensity (%MSO/×tradMT)
lower ankle point
38.9 %MSO
0.93 × tradMT
tradMT (100 µV)
42 %MSO
1 × tradMT
inflection point
44.9 %MSO
1.07 × tradMT
SI at 100 µV
45.2 %MSO
1.08 × tradMT
higher ankle point
50.8 %MSO
1.21 × tradMT

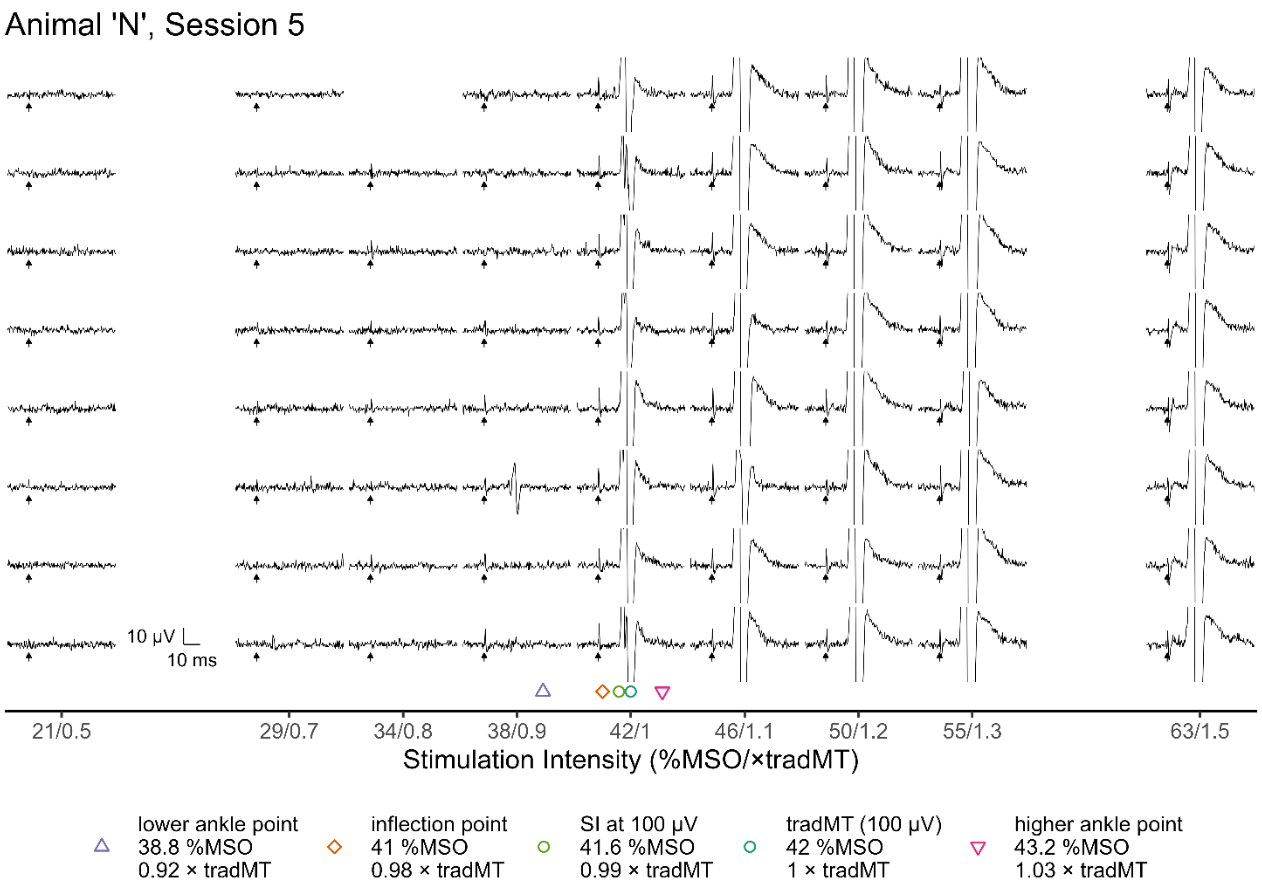
Animal 'N', Session 5
10 µV
10 ms
21/0.5
29/0.7
34/0.8
38/0.9
42/1
46/1.1
50/1.2
55/1.3
63/1.5
Stimulation Intensity (%MSO/×tradMT)
lower ankle point
38.8 %MSO
0.92 × tradMT
inflection point
41 %MSO
0.98 × tradMT
SI at 100 µV
41.6 %MSO
0.99 × tradMT
tradMT (100 µV)
42 %MSO
1 × tradMT
higher ankle point
43.2 %MSO
1.03 × tradMT

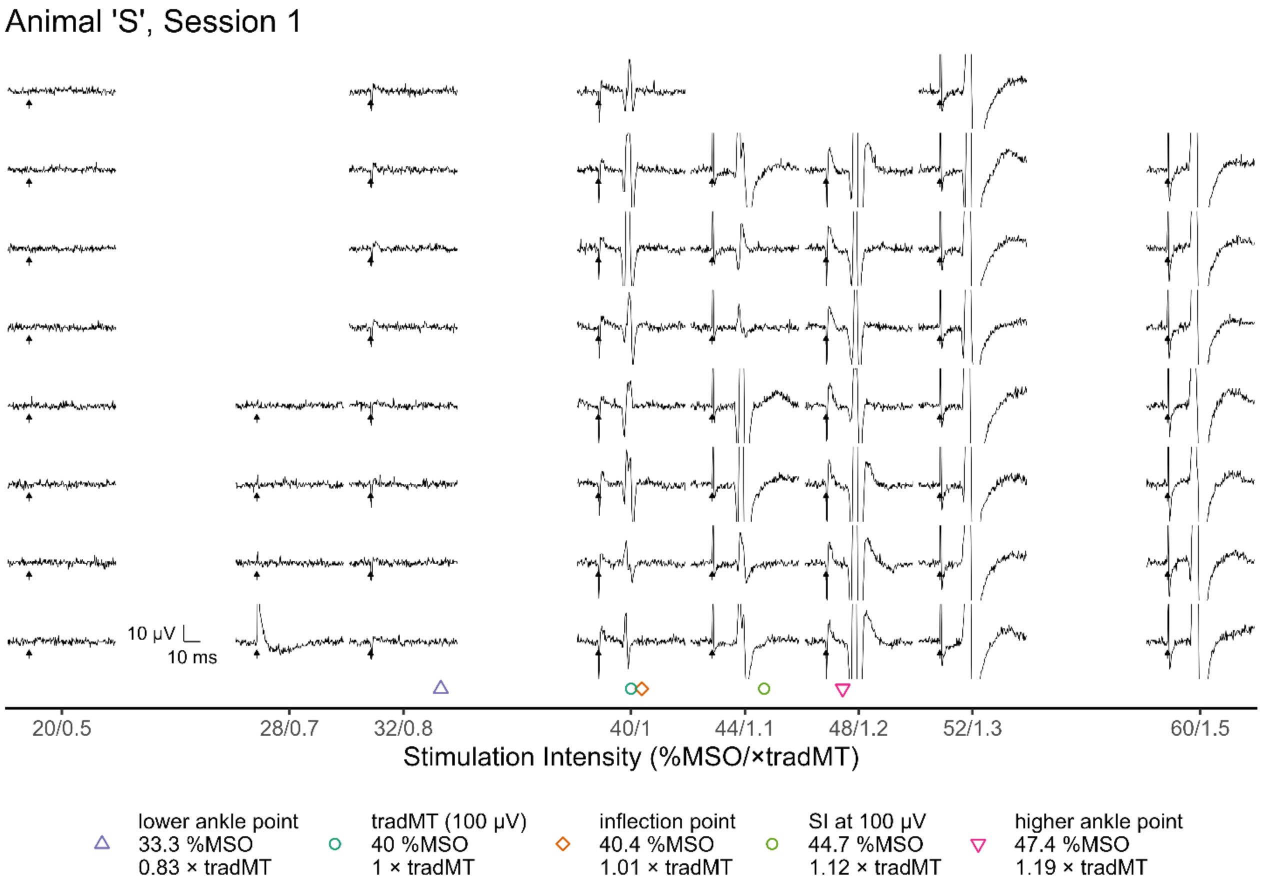
Animal 'S', Session 1
10 µV
10 ms
20/0.5
28/0.7
32/0.8
40/1
44/1.1
48/1.2
52/1.3
60/1.5
Stimulation Intensity (%MSO/×tradMT)
lower ankle point
33.3 %MSO
0.83 × tradMT
tradMT (100 µV)
40 %MSO
1 × tradMT
inflection point
40.4 %MSO
1.01 × tradMT
SI at 100 µV
44.7 %MSO
1.12 × tradMT
higher ankle point
47.4 %MSO
1.19 × tradMT

Animal 'S', Session 2

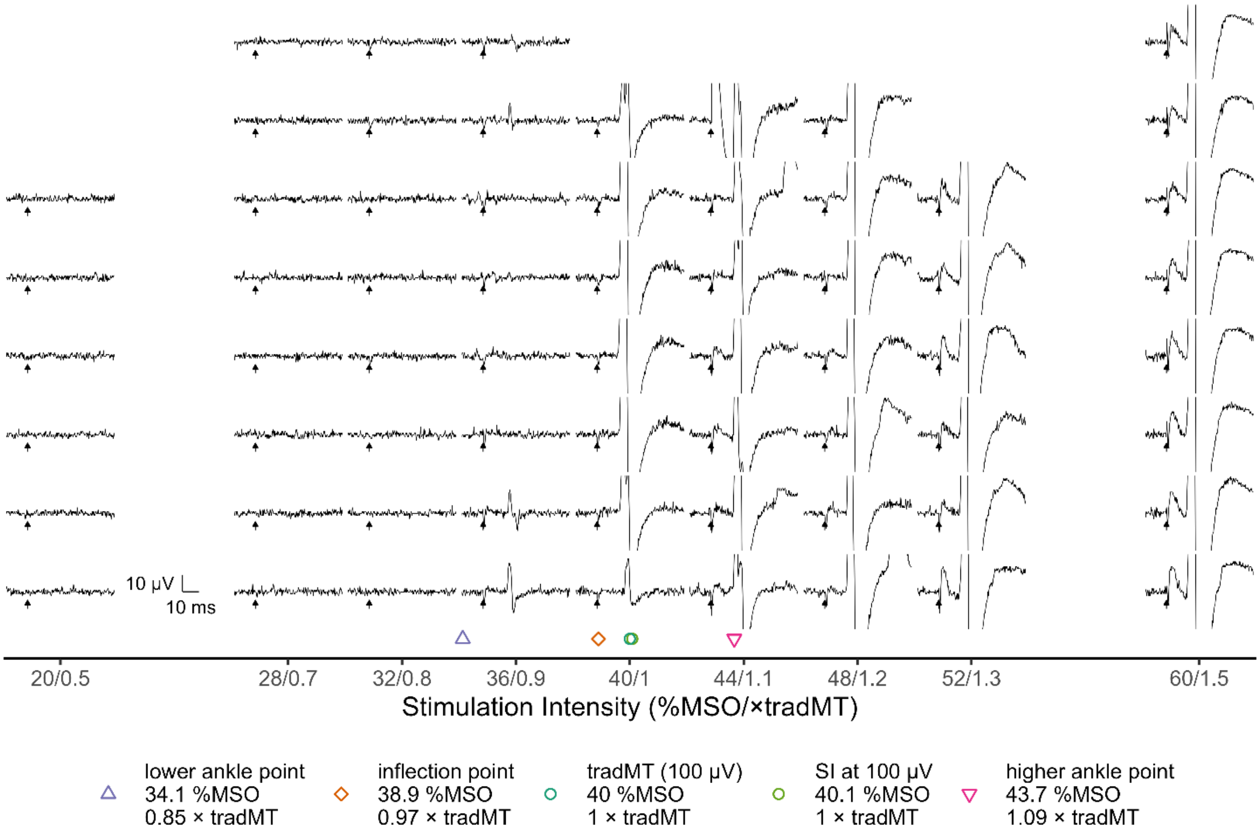


Animal 'S', Session 3

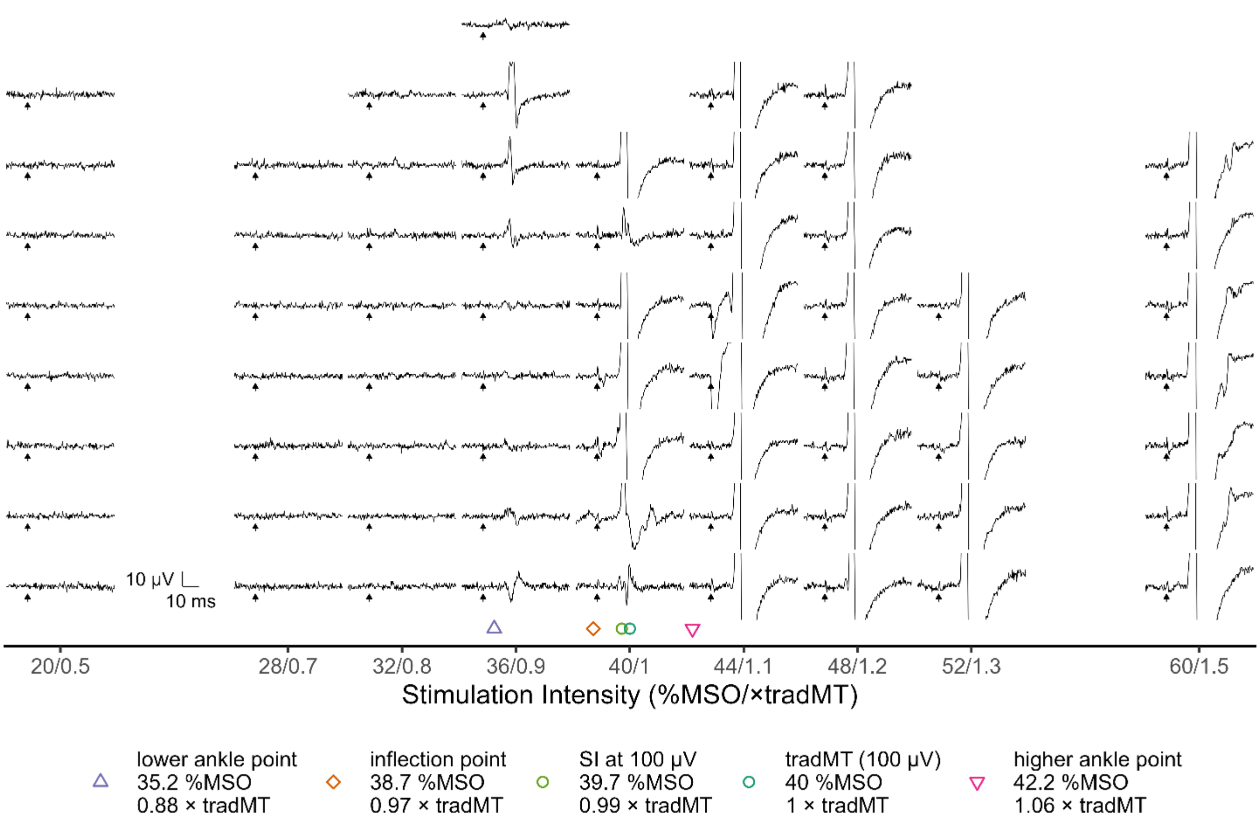

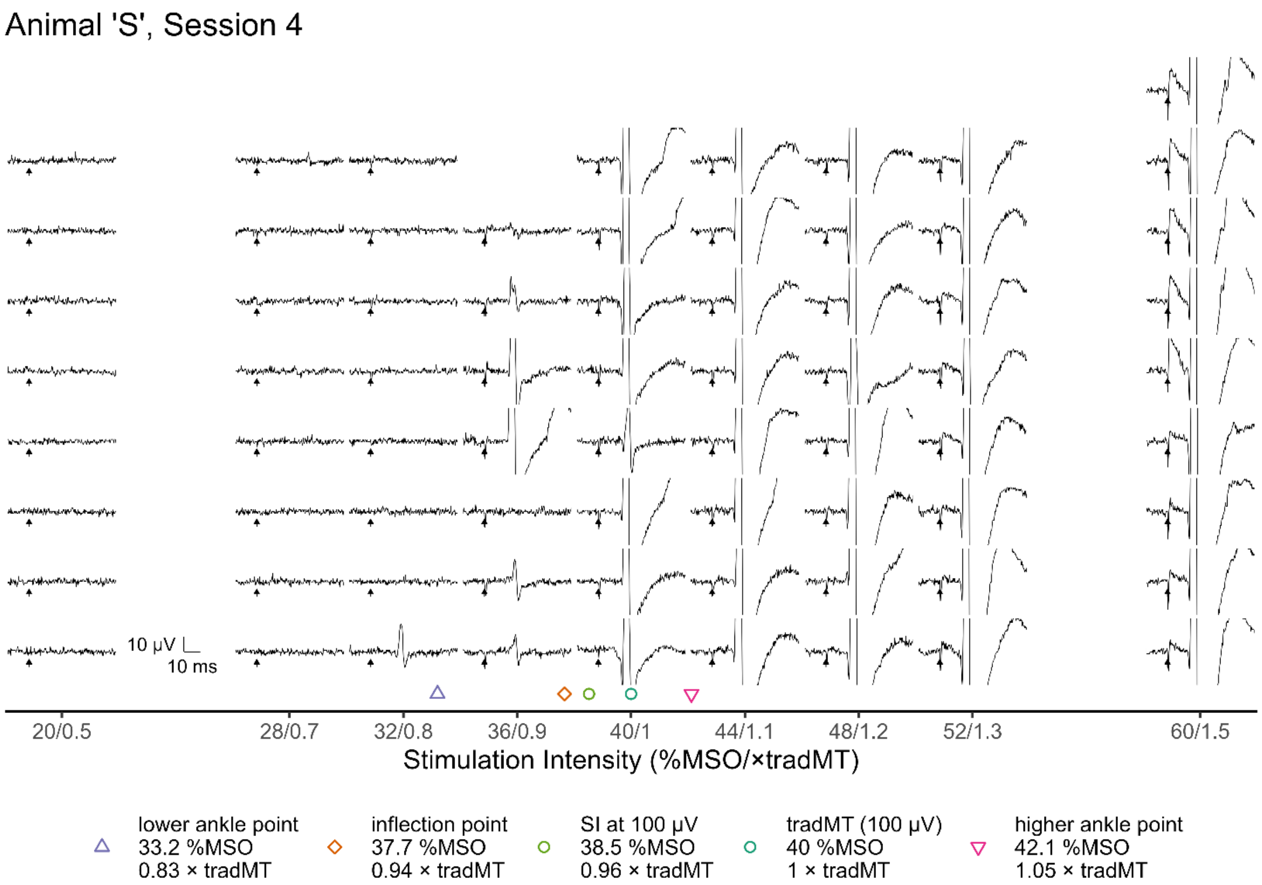
Animal 'S', Session 4
10 µV
10 ms
20/0.5
28/0.7
32/0.8
36/0.9
40/1
44/1.1
48/1.2
52/1.3
60/1.5
Stimulation Intensity (%MSO/×tradMT)
lower ankle point 33.2 %MSO 0.83 × tradMT
inflection point 37.7 %MSO 0.94 × tradMT
SI at 100 µV 38.5 %MSO 0.96 × tradMT
tradMT (100 µV) 40 %MSO 1 × tradMT
higher ankle point 42.1 %MSO 1.05 × tradMT

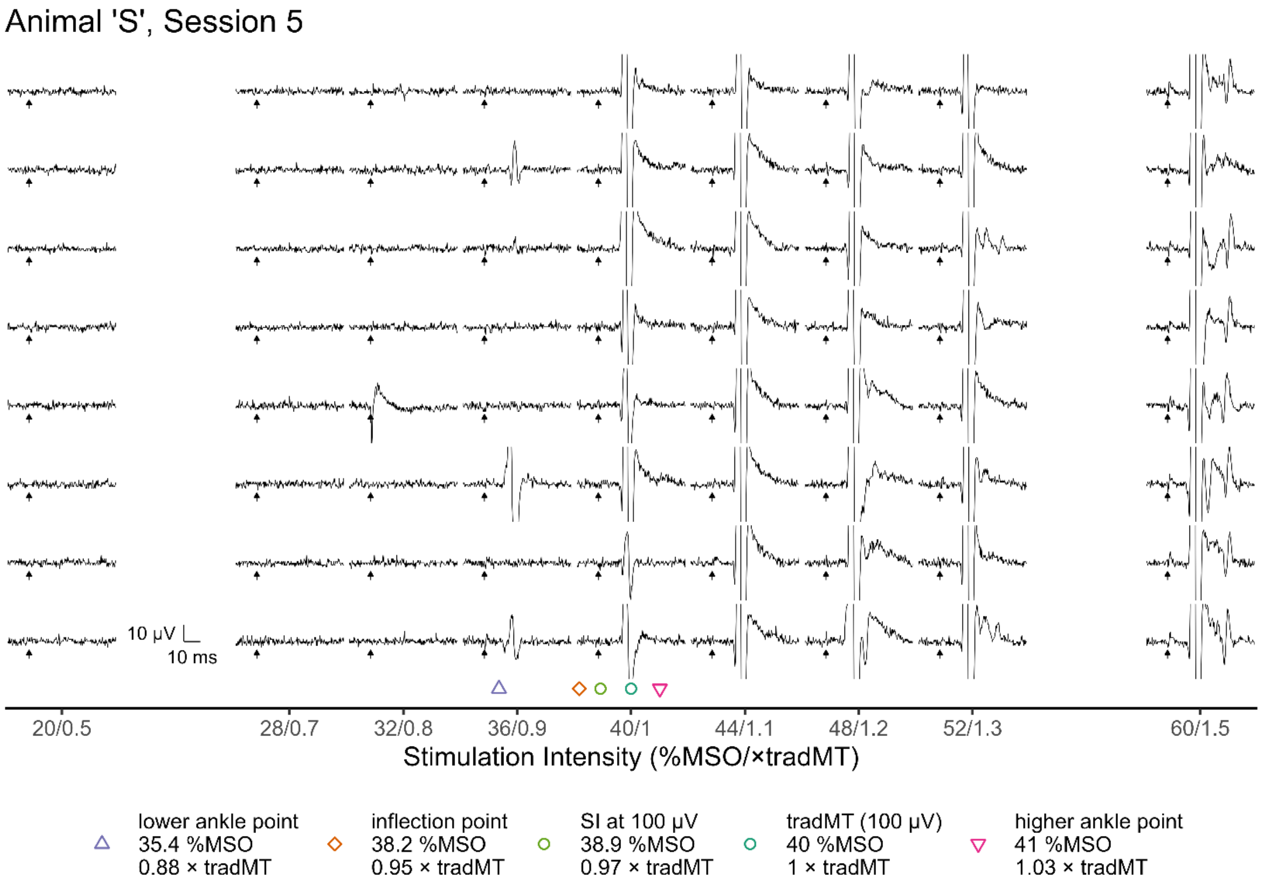
Animal 'S', Session 5
10 µV
10 ms
20/0.5
28/0.7
32/0.8
36/0.9
40/1
44/1.1
48/1.2
52/1.3
60/1.5
Stimulation Intensity (%MSO/×tradMT)
lower ankle point 35.4 %MSO 0.88 × tradMT
inflection point 38.2 %MSO 0.95 × tradMT
SI at 100 µV 38.9 %MSO 0.97 × tradMT
tradMT (100 µV) 40 %MSO 1 × tradMT
higher ankle point 41 %MSO 1.03 × tradMT

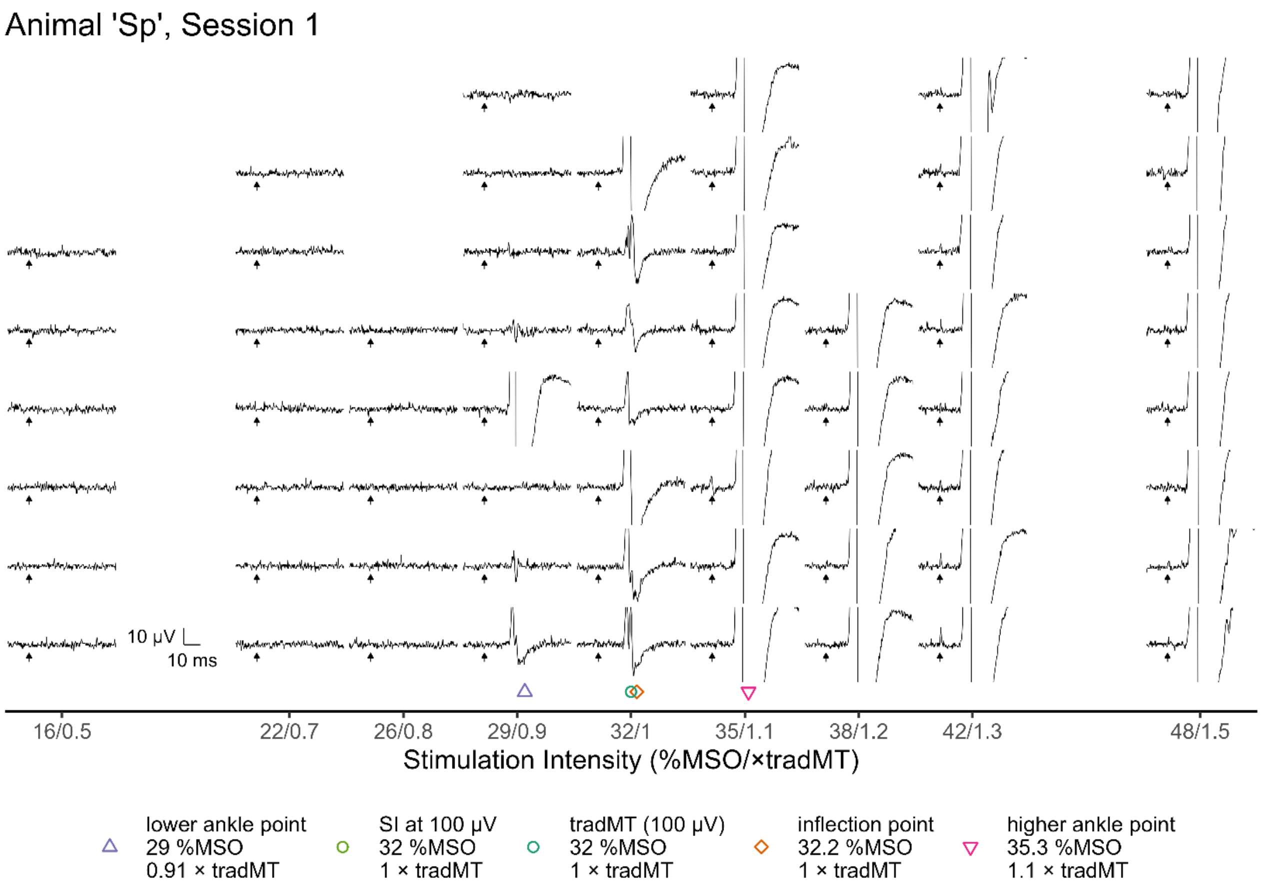
Animal 'Sp', Session 1
10 µV
10 ms
16/0.5
22/0.7
26/0.8
29/0.9
32/1
35/1.1
38/1.2
42/1.3
48/1.5
Stimulation Intensity (%MSO/×tradMT)
lower ankle point
29 %MSO
0.91 × tradMT
SI at 100 µV
32 %MSO
1 × tradMT
tradMT (100 µV)
32 %MSO
1 × tradMT
inflection point
32.2 %MSO
1 × tradMT
higher ankle point
35.3 %MSO
1.1 × tradMT

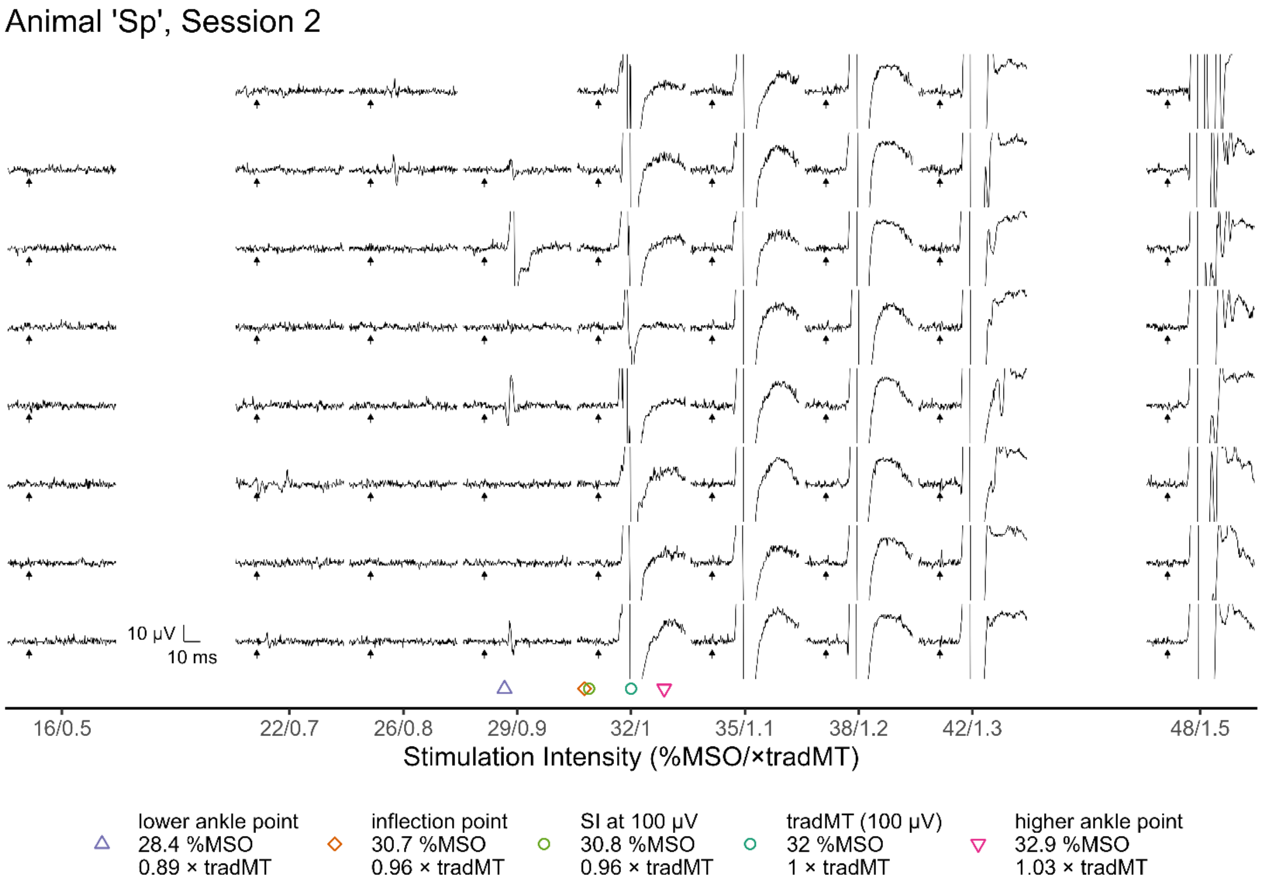
Animal 'Sp', Session 2
10 µV
10 ms
16/0.5
22/0.7
26/0.8
29/0.9
32/1
35/1.1
38/1.2
42/1.3
48/1.5
Stimulation Intensity (%MSO/×tradMT)
lower ankle point
28.4 %MSO
0.89 × tradMT
inflection point
30.7 %MSO
0.96 × tradMT
SI at 100 µV
30.8 %MSO
0.96 × tradMT
tradMT (100 µV)
32 %MSO
1 × tradMT
higher ankle point
32.9 %MSO
1.03 × tradMT

Animal 'Sp', Session 3

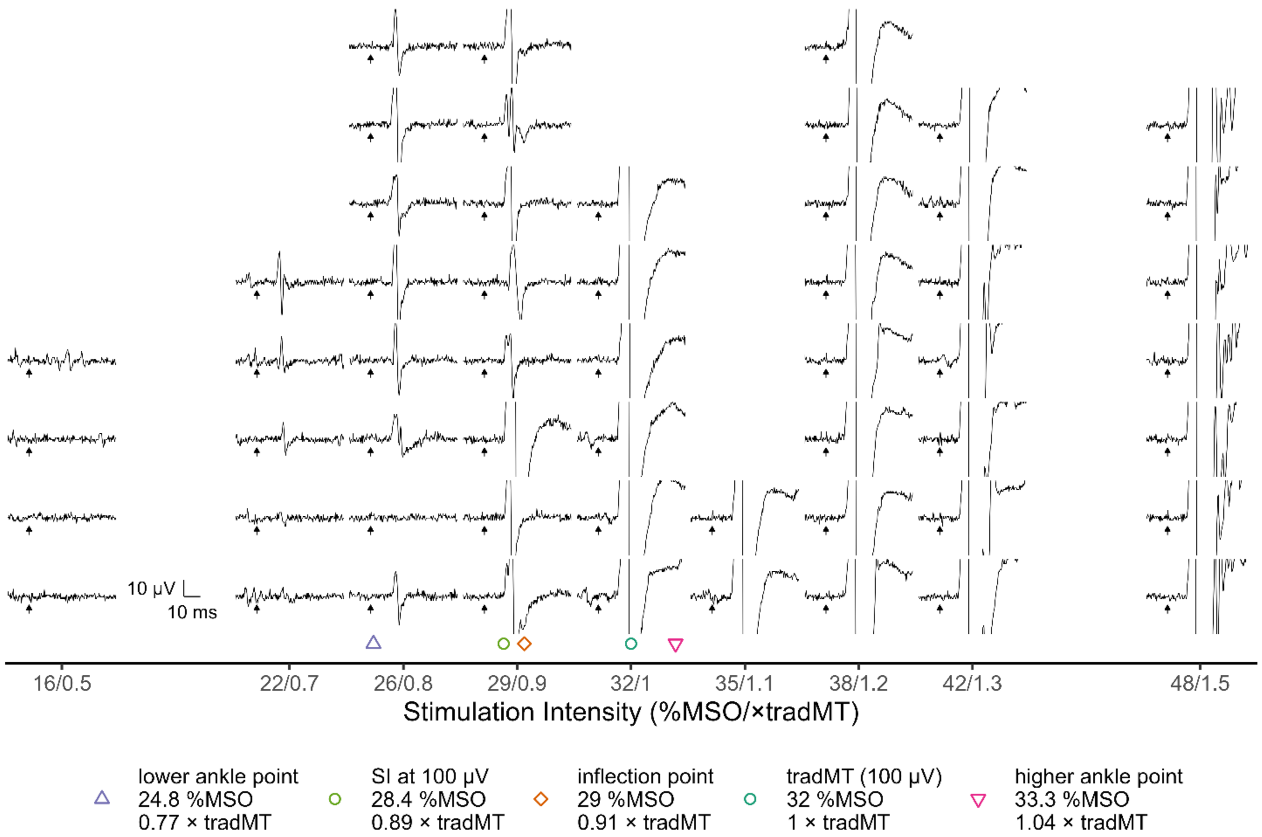


Animal 'Sp', Session 4

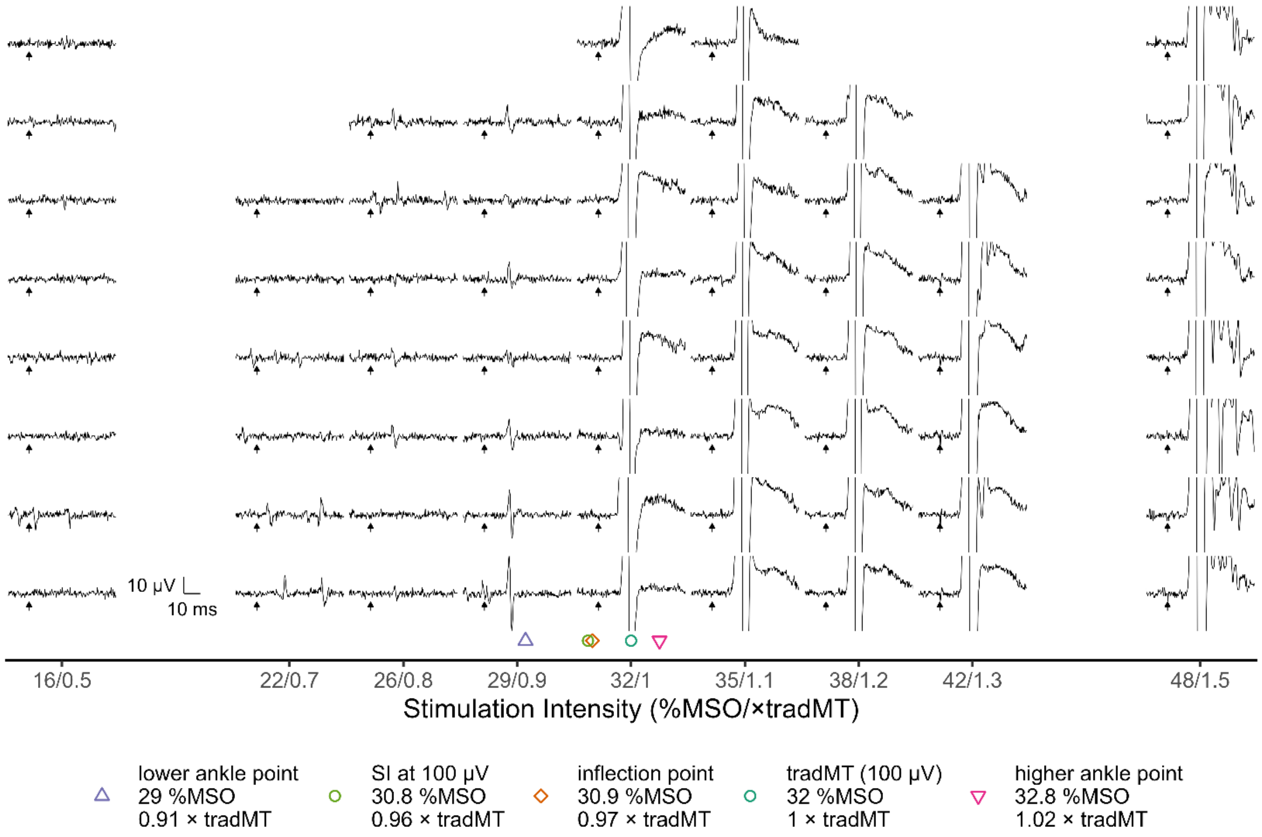

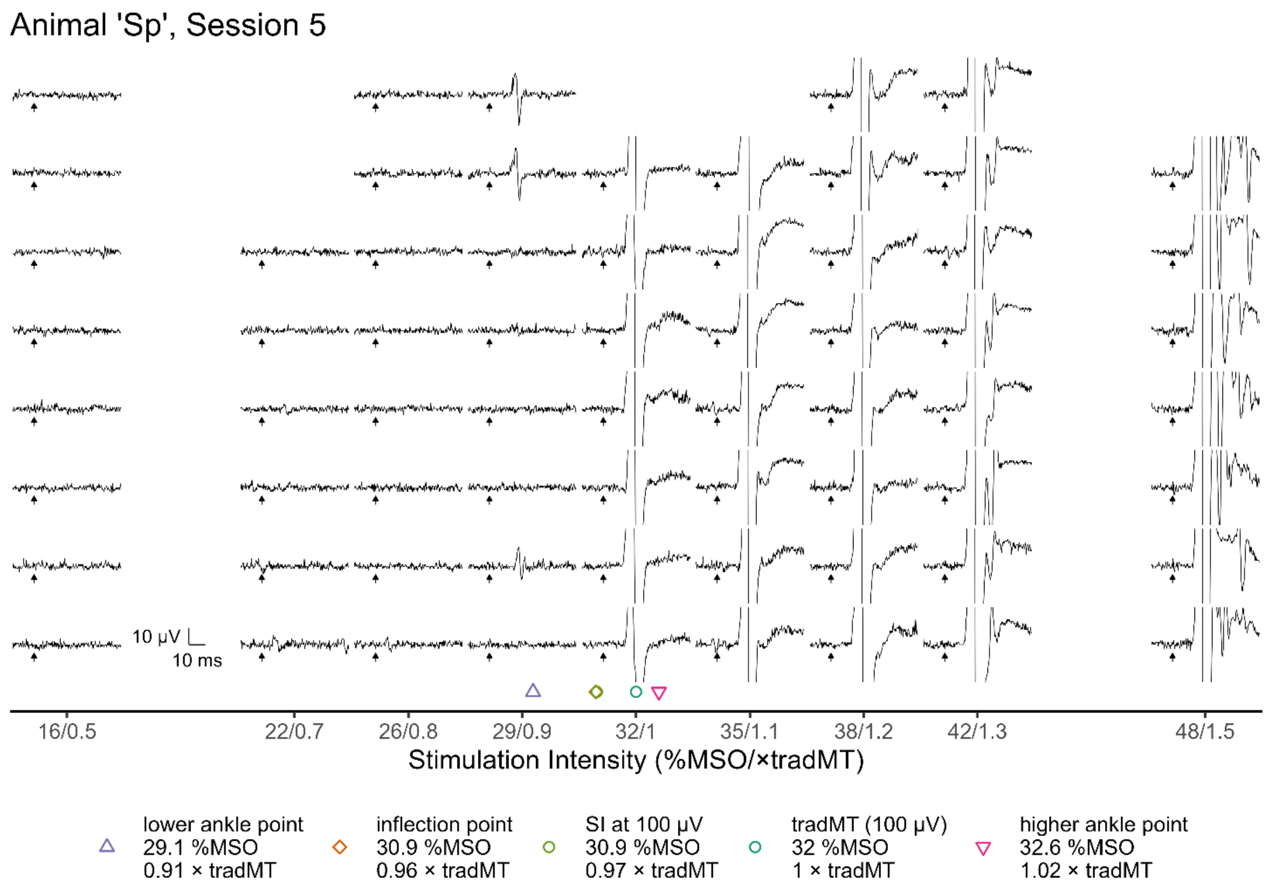
Animal 'Sp', Session 5
10 μV
10 ms
16/0.5
22/0.7
26/0.8
29/0.9
32/1
35/1.1
38/1.2
42/1.3
48/1.5
Stimulation Intensity (%MSO/×tradMT)
lower ankle point 29.1 %MSO 0.91 × tradMT
inflection point 30.9 %MSO 0.96 × tradMT
SI at 100 μV 30.9 %MSO 0.97 × tradMT
tradMT (100 μV) 32 %MSO 1 × tradMT
higher ankle point 32.6 %MSO 1.02 × tradMT

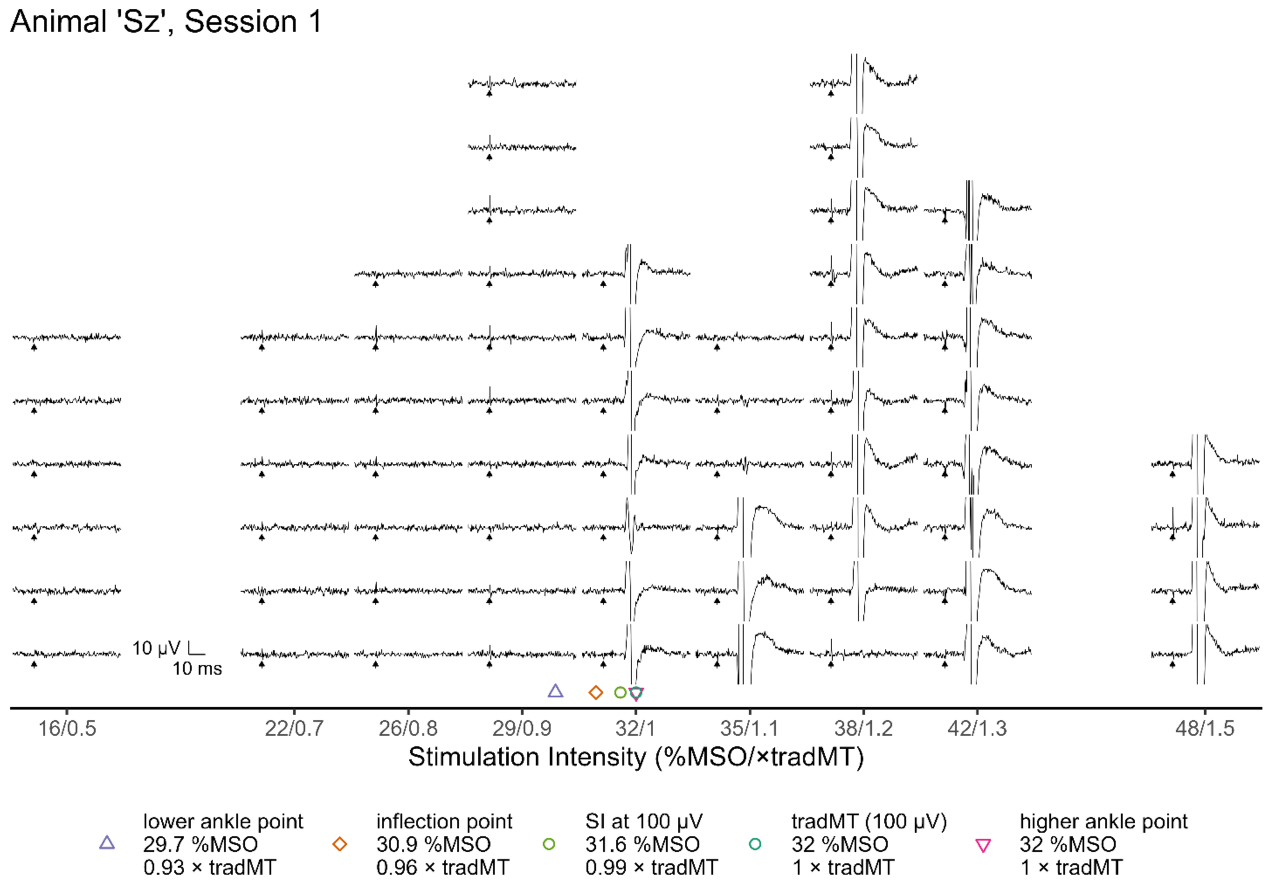
Animal 'Sz', Session 1
10 μV
10 ms
16/0.5
22/0.7
26/0.8
29/0.9
32/1
35/1.1
38/1.2
42/1.3
48/1.5
Stimulation Intensity (%MSO/×tradMT)
lower ankle point 29.7 %MSO 0.93 × tradMT
inflection point 30.9 %MSO 0.96 × tradMT
SI at 100 μV 31.6 %MSO 0.99 × tradMT
tradMT (100 μV) 32 %MSO 1 × tradMT
higher ankle point 32 %MSO 1 × tradMT

Animal 'Sz', Session 2

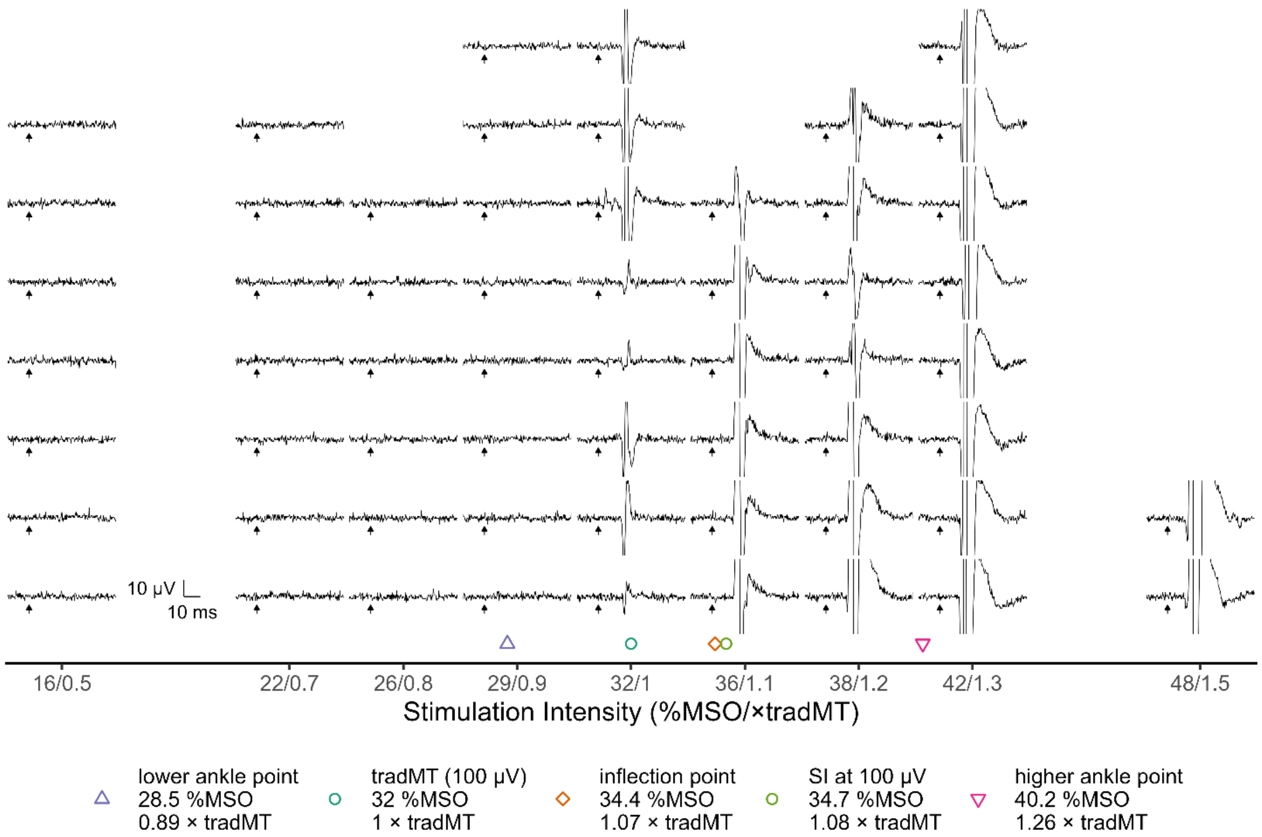


Animal 'Sz', Session 3

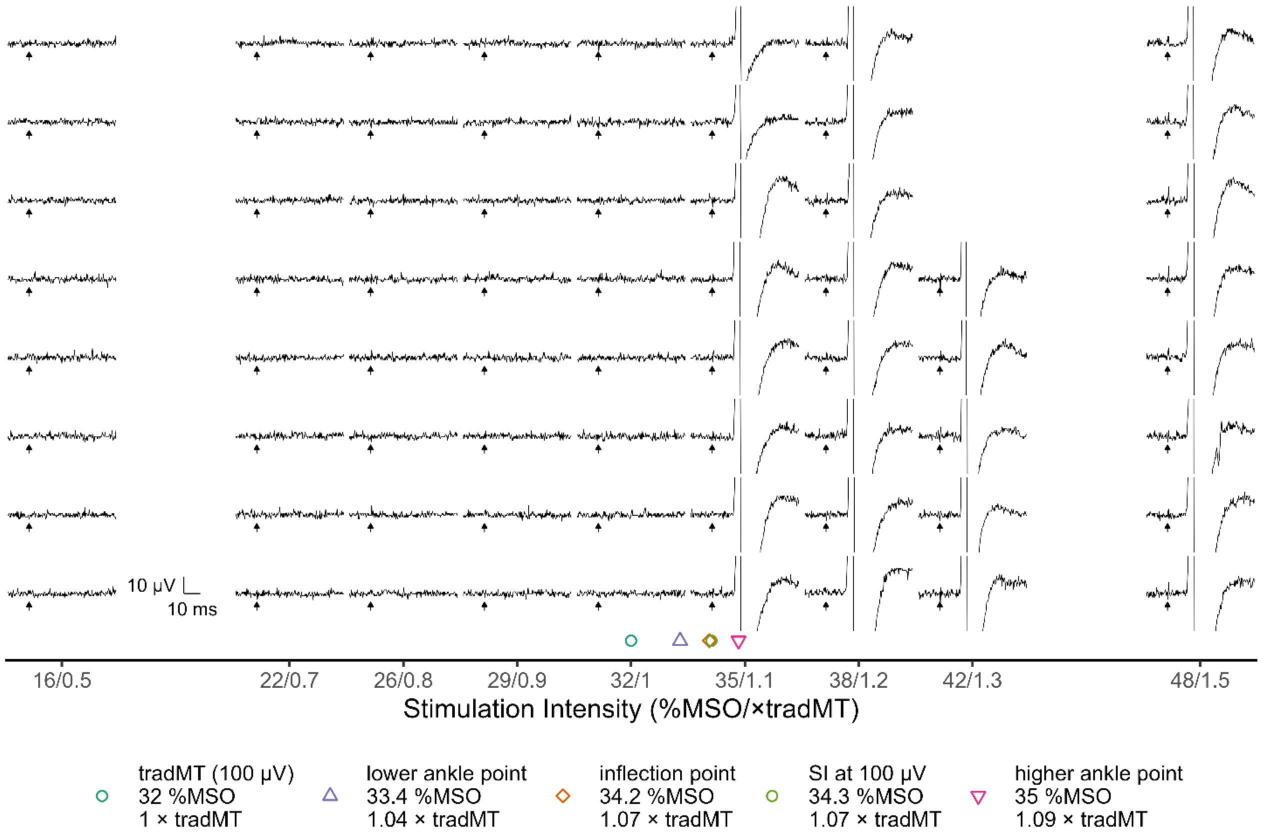

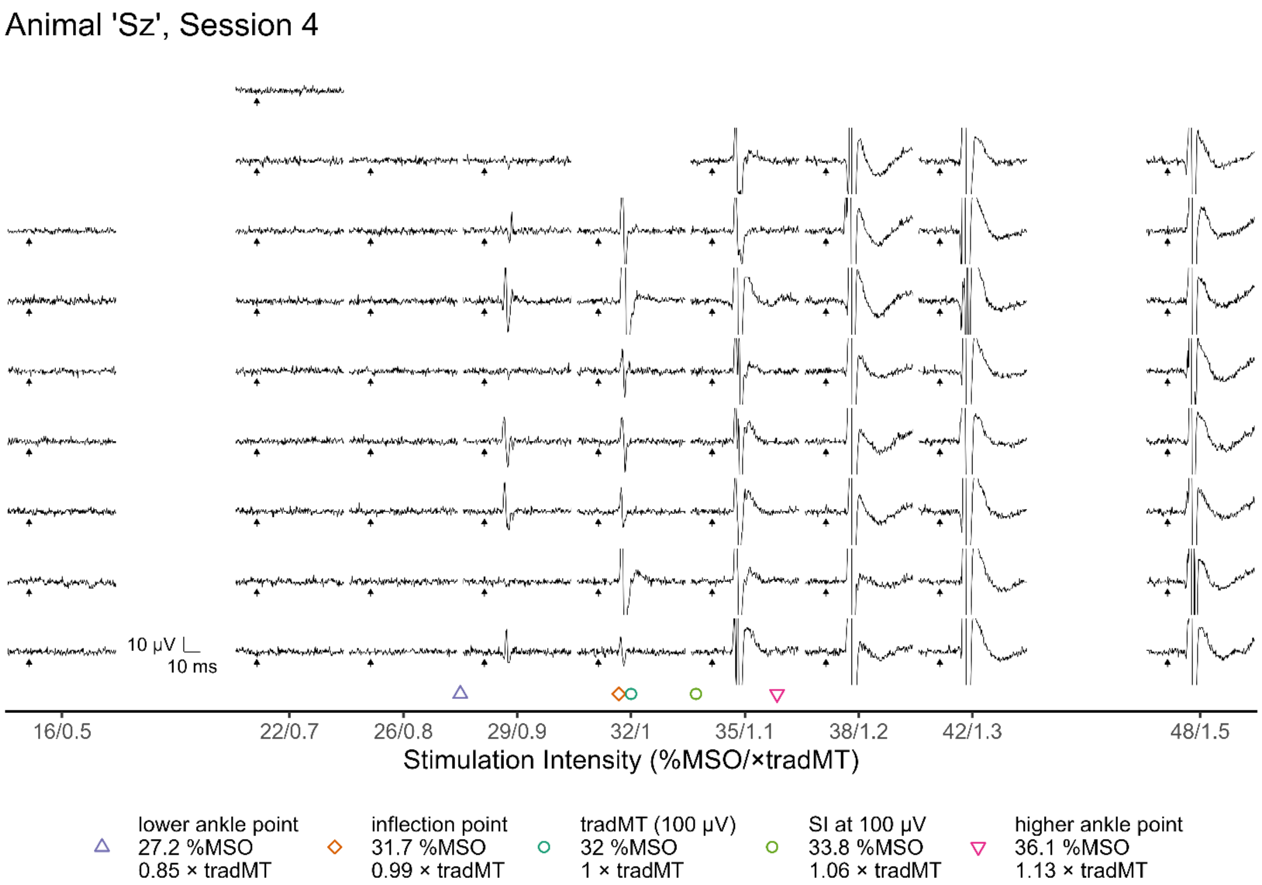
Animal 'Sz', Session 4
10 µV
10 ms
16/0.5
22/0.7
26/0.8
29/0.9
32/1
35/1.1
38/1.2
42/1.3
48/1.5
Stimulation Intensity (%MSO/×tradMT)
lower ankle point
27.2 %MSO
0.85 × tradMT
inflection point
31.7 %MSO
0.99 × tradMT
tradMT (100 µV)
32 %MSO
1 × tradMT
SI at 100 µV
33.8 %MSO
1.06 × tradMT
higher ankle point
36.1 %MSO
1.13 × tradMT

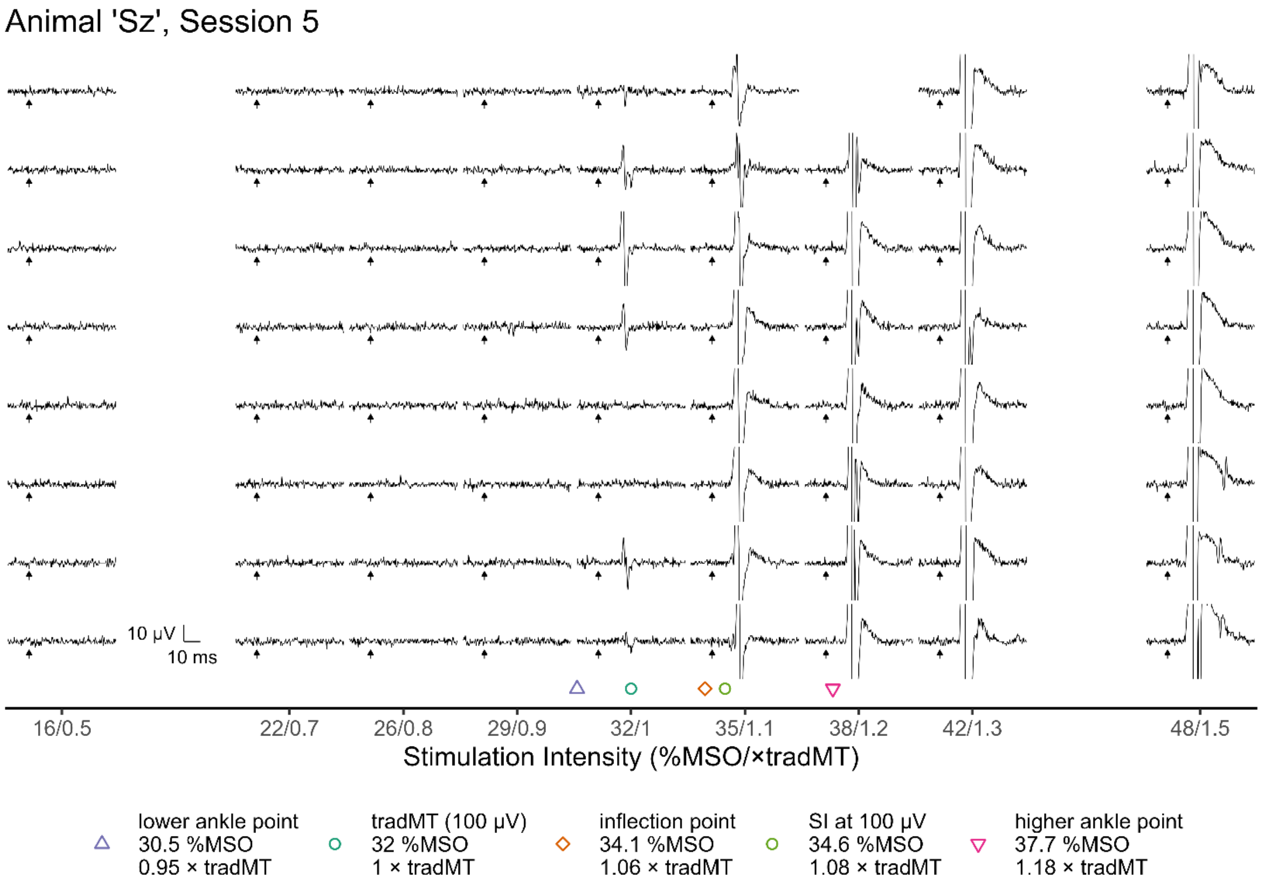
Animal 'Sz', Session 5
10 µV
10 ms
16/0.5
22/0.7
26/0.8
29/0.9
32/1
35/1.1
38/1.2
42/1.3
48/1.5
Stimulation Intensity (%MSO/×tradMT)
lower ankle point
30.5 %MSO
0.95 × tradMT
tradMT (100 µV)
32 %MSO
1 × tradMT
inflection point
34.1 %MSO
1.06 × tradMT
SI at 100 µV
34.6 %MSO
1.08 × tradMT
higher ankle point
37.7 %MSO
1.18 × tradMT
